# Modulation of Biophysical Properties of Nucleocapsid Protein in the Mutant Spectrum of SARS-CoV-2

**DOI:** 10.1101/2023.11.21.568093

**Authors:** Ai Nguyen, Huaying Zhao, Dulguun Myagmarsuren, Sanjana Srinivasan, Di Wu, Jiji Chen, Grzegorz Piszczek, Peter Schuck

## Abstract

Genetic diversity is a hallmark of RNA viruses and the basis for their evolutionary success. Taking advantage of the uniquely large genomic database of SARS-CoV-2, we examine the impact of mutations across the spectrum of viable amino acid sequences on the biophysical phenotypes of the highly expressed and multifunctional nucleocapsid protein. We find variation in the physicochemical parameters of its extended intrinsically disordered regions (IDRs) sufficient to allow local plasticity, but also exhibiting functional constraints that similarly occur in related coronaviruses. In biophysical experiments with several N-protein species carrying mutations associated with major variants, we find that point mutations in the IDRs can have nonlocal impact and modulate thermodynamic stability, secondary structure, protein oligomeric state, particle formation, and liquid-liquid phase separation. In the Omicron variant, distant mutations in different IDRs have compensatory effects in shifting a delicate balance of interactions controlling protein assembly properties, and include the creation of a new protein-protein interaction interface in the N-terminal IDR through the defining P13L mutation. A picture emerges where genetic diversity is accompanied by significant variation in biophysical characteristics of functional N-protein species, in particular in the IDRs.

## Introduction

A salient characteristic of RNA viruses is their high error rate in transcription and their resulting quasispecies nature (Eigen, 1996; Holland and Domingo, 1997). This diversity is also reflected in the ensemble of consensus sequences sampled across the infected host population, as is apparent in the GISAID (Global Initiative on Sharing All Influenza Data) repository of SARS-CoV-2 genomes (Elbe and Buckland-Merrett, 2017). With currently ≈15 million entries, this unprecedented large database has provided the basis for phylogenetic analyses that have identified critical amino acid mutations associated with immune evasion, infectivity, and disease severity, and allowed the rapid identification of variants of concern (Greaney et al., 2022; Kepler et al., 2021; Obermeyer et al., 2022; Rochman et al., 2021; Viana et al., 2022). The vast majority of mutations, however, seem inconsequential in that they usually do not lead to any fixed substitutions. Nonetheless, the mutant spectrum exhaustively describes a landscape of amino acids that may occupy any position in the viral proteins, as in a natural deep mutational scan (Bloom and Neher, 2023; Schuck and Zhao, 2023; Zhao et al., 2022). Biophysical constraints implicit in the shape of such landscapes are key to understand the function and molecular evolution of viral proteins (Starr and Thornton, 2016; Wang et al., 2021).

Unfortunately, the wealth of genomic information on SARS-CoV-2 stands in stark contrast with our knowledge of the phenotypic consequences of sequence mutations. In conjunction with biophysical and structural studies, inspections of local mutations have increased our understanding of mechanisms of SARS-CoV-2 entry, mechanisms of replication and assembly, and interaction with various host factors (Dadonaite et al., 2023; Del Veliz et al., 2021; Greaney et al., 2022; Hu et al., 2023; Stevens et al., 2022; Syed et al., 2021; Zhao et al., 2023, 2022). Furthermore, the range of naturally occurring mutations at target sites is an important consideration for potential drugs, vaccines, and diagnostics (Artesi et al., 2020; Saldivar-Espinoza et al., 2022; Tian et al., 2022). Outside these focused studies of relatively well-understood hot spots, however, the mutational landscape has remained relatively unexplored.

Biophysical fitness landscapes have been studied with regard to observables such as thermal stability of globular proteins, solvent accessibility, catalytic activity, or binding affinity of protein-protein interfaces, which has led to significant advances in understanding relationship between molecular properties, population fitness, and evolutionary processes (Bershtein et al., 2017; Bloom et al., 2006; Echave and Wilke, 2017; Lässig et al., 2017; Liberles et al., 2012; Serohijos and Shakhnovich, 2014; Sikosek and Chan, 2014; Wang et al., 2015). However, it was found that constraints for evolution of intrinsically disordered regions (IDRs) are much different from those of globular proteins (Brown et al., 2010; Lafforgue et al., 2022). Generally, intrinsic disorder and loose packing is a common characteristic of many RNA virus proteins (Tokuriki et al., 2009), which is thought to promote functional promiscuity, permit greater diversity, and enhance evolvability to adopt new functions with few mutations (Charon et al., 2018; Gitlin et al., 2014; Tokuriki and Tawfik, 2009). One possible mechanism is viral mimicry of host-protein short linear motifs (SLiMs) that allow binding to host protein domains and cause subversion of host cellular pathways (Davey et al., 2015, 2011; Hagai et al., 2014; Kruse et al., 2021; Mihalič et al., 2023; Schuck and Zhao, 2023; Shuler and Hagai, 2022). It was also shown how nonlocal biophysical properties, such as the charge of intrinsically disordered regions (IDRs), can be relevant evolutionary traits (Zarin et al., 2021, 2017). More recently, it was recognized that the formation of membrane-less cellular compartments driven by liquid-liquid phase separation (LLPS) is a key aspect of many intrinsically disordered proteins, including many viral proteins (Cascarina and Ross, 2022; Zhang et al., 2023). What kind of sequence constraints may derive from the biophysical requirement to conserve LLPS properties is currently only emerging (Brown et al., 2011; Chin et al., 2022; Ho and Huang, 2022; Lin et al., 2017; Riback et al., 2017).

The goal of the present work is to probe the phenotypic diversity with respect to several biophysical properties of SARS-CoV-2 nucleocapsid (N-)protein, taking advantage of the vast mutational landscape of SARS-CoV-2. N-protein is the most abundant viral protein in the infected cell (Finkel et al., 2021), and as we reported previously (Zhao et al., 2022), it is also the most diverse structural protein with approximately 86% of its 419 residues capable of assuming on average 4 to 5 different amino acids evidently without impairment of viability. The highest frequency of mutations occurs in the substantial IDRs which are the N-arm, linker, and C-arm that flank and connect the folded nucleic acid binding domain (NTD) and the dimerization domain (CTD) (**Figure 1**). The IDRs comprise approximately half of the molecule and allow large conformational fluctuations (Botova et al., 2024; Cubuk et al., 2021; Redzic et al., 2021). The eponymous structural function of N-protein is that of scaffolding genomic RNA for virion assembly. It proceeds *via* nucleic acid (NA)-binding induced conformational changes and oligomerization, leading to the formation of ribonucleoprotein (RNP) particles with as-of-yet unknown molecular architecture, ≈38 of which are arranged like beads-on-a-string in the viral particle (Carlson et al., 2022; Cubuk et al., 2021; Klein et al., 2020; Yao et al., 2020; Zhao et al., 2024, 2023, 2021), and are anchored through binding of N-protein to viral M-protein (Lu et al., 2021; Masters, 2019). Beyond this structural role, N-protein is highly multi-functional and binds to multiple host proteins to modulate or exploit different pathways, including stress granules (Biswal et al., 2022; Gordon et al., 2020; Savastano et al., 2020), the type 1 interferon signaling pathway (Chen et al., 2020; Li et al., 2020), the NLRP3 inflammasome (Pan et al., 2021), and others, as recently reviewed (Wu et al., 2023; Yu et al., 2023). N-protein can form macromolecular condensates through liquid-liquid phase separation (LLPS) that aid in assembly functions and interactions with host proteins (Carlson et al., 2020; Cascarina and Ross, 2022; Cubuk et al., 2021; Iserman et al., 2020; Jack et al., 2021; Lu et al., 2021; Perdikari et al., 2020; Savastano et al., 2020). In addition, it is also localized at exterior cell surfaces, where it was found to bind many different chemokines, likely manipulating innate immunity through chemokine sequestration (López-Muñoz et al., 2022).

**Figure 1.**
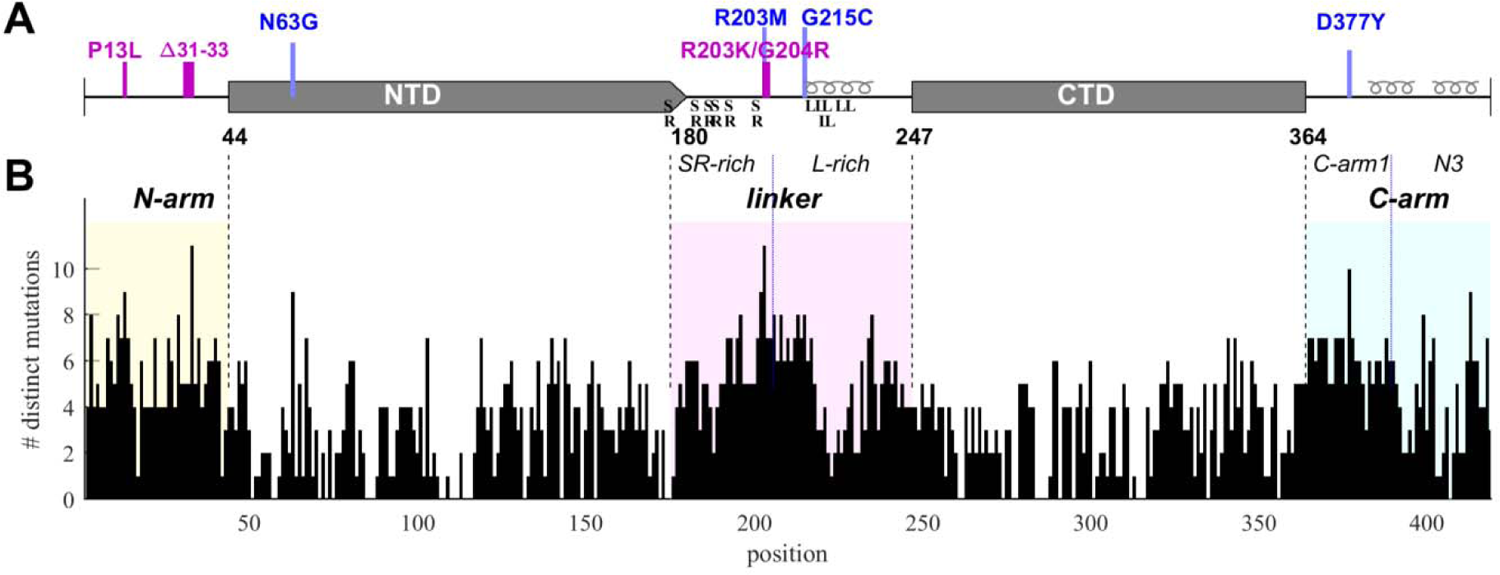
Structural organization and sequence plasticity of N-protein. (A) Schematics of folded regions (NTD and CTD, rectangles) and disordered regions (N-arm, linker, and C-arm, straight line) along the N-protein sequence. Defining mutations from the Delta-variant are indicated in blue, those from Omicron-variants in magenta. Transient helices in the disordered regions are highlighted, as well as SR-rich and L-rich linker sequences and the C-terminal N3 region. (B) Histogram of the number of distinct amino acid mutations at each position. For clarity and reference to other figures, IDRs are shaded with N-arm highlighted in yellow, linker in magenta, and C-arm in cyan.

The large number of structural and non-structural N-protein functions poses the question of how they are conserved in light of the significant sequence diversity. In the present work we computationally evaluate the range of several biophysical traits resulting from diversity in the SARS-CoV-2 N-protein folded domains and IDRs across the observed mutant spectrum, as well as related coronaviruses. In complementary biophysical experiments with several representative N-protein mutants derived from SARS-CoV-2 variants of concern we characterize their variation in thermodynamic stability, secondary structure, oligomeric state, energetics of NA binding, assembly and LLPS propensity. We find that a large biophysical parameter space is available for viable N-protein, with the potential for mutations to exert nonlocal effects modulating overall protein biophysical properties.

## Results

### Distribution of physicochemical properties across the SARS-CoV-2 mutant spectrum

SARS-CoV-2 sequence data were downloaded from Nextstrain (Hadfield et al., 2018) in January 2023 and 5.06 million high quality sequences were selected for analysis. The N-protein amino acid sequences exhibit ≈43 million instances of mutations distributed across ≈92% of its residues. We have previously characterized this dataset with regard to the amino acid mutational landscape of N-protein, and found mutation frequencies that are strongly dependent on position and largely time-invariant, except for the defining mutations arising in variants of concern, the latter comprising ≈36% Delta-variant and ≈49% Omicron-variant sequences (Schuck and Zhao, 2023). A histogram of the number of different amino acids mutations that are found at each residue is shown in **Figure 1B**. It may be discerned that sequence plasticity is highest in the IDRs, with an average of 5.2 different possible amino acid mutations at each residue compared to 2.9 different mutations on average in the folded domains.

Exploiting the N-protein mutational landscape and sequence data, previous work in our laboratory has focused on local amino acid sequence properties such as mutation effects on transient structural features in the linker IDR (Zhao et al., 2023) and the creation of short linear motifs (Schuck and Zhao, 2023). However, nonlocal biophysical properties may also be functionally critical and evolutionarily conserved despite amino acid sequence heterogeneity in IDRs (Zarin et al., 2021, 2017). The sequence ensembles extracted from the genomic database allow us to ask whether physicochemical properties are constrained or can vary across viable sequences of the mutation spectrum.

To this end, genome data were sorted into unique groups with distinct N-protein amino acid sequences, each sequence carrying a set of distinct mutations that represent a viable N-protein species. For a robust analysis, each mutated sequence was required to be represented in at least 10 different genomes in the database. This led to 6,300 distinct full-length N-protein sequences (N-FL; 1-419). We similarly subdivided the N-protein in different regions (**Figure 1A**) and grouped unique sets of mutations in each region: For the folded domains we found 720 distinct NTD (N:45-179) and 399 distinct CTD (N:248-363) sequences, while for the IDRs there are 512 N-arm (N:1-44), 1039 linker (N:175-247), and 556 C-arm (N:364-419) sequences. (Due to ambiguity in delineation between NTD and linker, designations overlapping in 175-180 were used to avoid artificial truncation and permit conservative evaluation of the properties of each domain.) Further subdividing the linker there are 349 distinct sequences for the SR-rich region (N:175-205) and 442 for the L-rich region (N:206-247), respectively. Finally, similarly subdividing the C-arm we obtained the 176 sequences for the N3 region (N:390-419) and 242 for the remainder of the C-arm (N:364-389).

We first examine polarity and hydrophobicity of N-protein and different regions based on their amino acid compositions. As shown in beehive plots of **Figure 2**, where each of the partially overlapping black dots represents one species from the cloud of mutant sequences, the index values of all N-FL sequences fall within a very narrow range (left column). Properties of the full-length protein may obscure significant differences on a smaller scale, in particular since the polarity and hydrophobicity indices are weighted-average properties. Focusing on folded N-protein modules, we find that hydrophobicity is uniformly high and polarity correspondingly low in the folded NTD and CTD domains, which is consistent with the expectation that folded structures are stabilized by buried hydrophobic residues (Eisenberg and McLachlan, 1986; Kauzmann, 1959). By contrast, IDRs exhibit significantly higher polarity and lower hydrophobicity. In particular, the N-arm and C-arm are most polar: despite a very large dispersion across the mutant spectrum, their values do not overlap with those of the folded domains.

**Figure 2.**
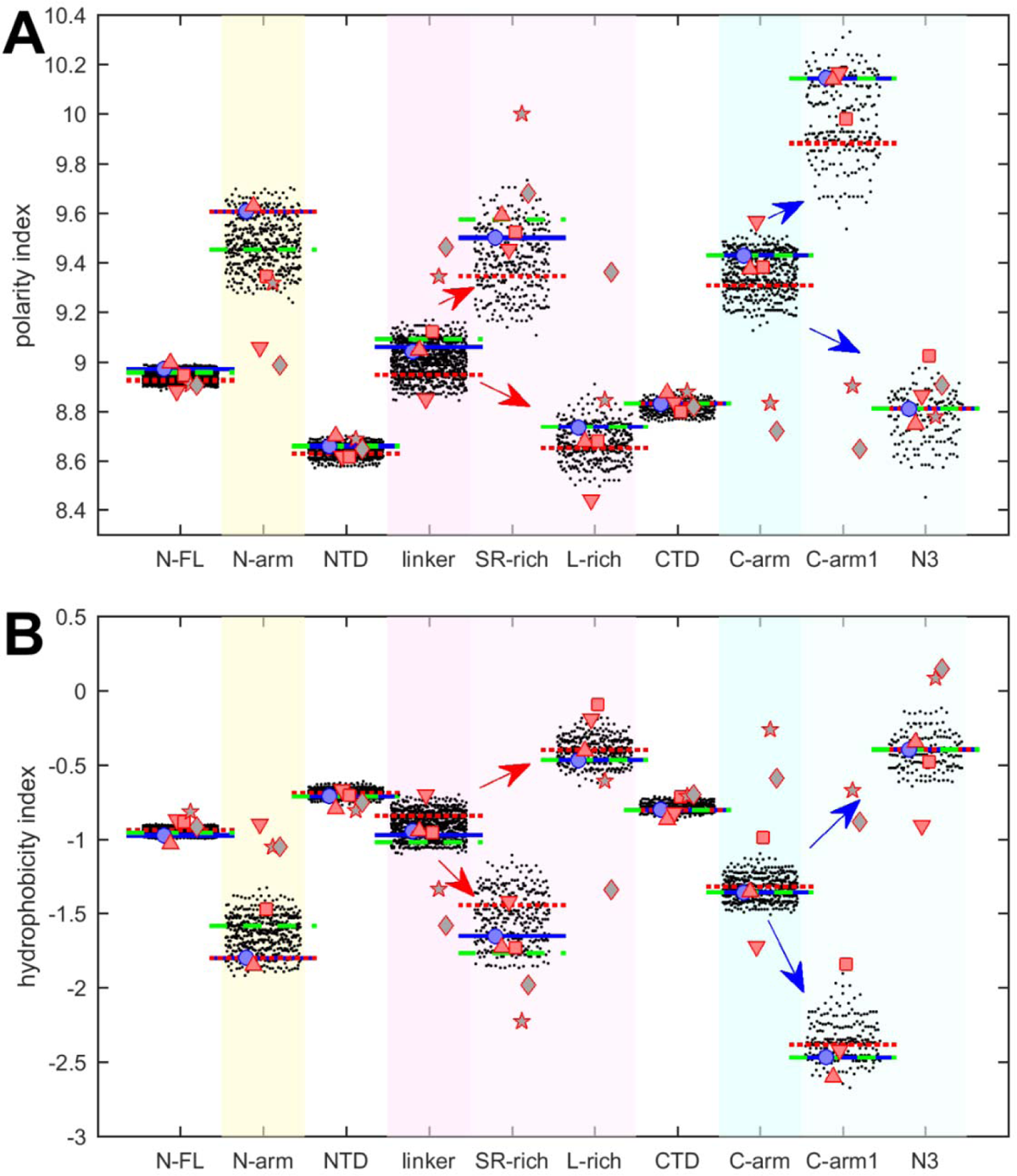
Beehive plots showing the distributions of polarity and hydrophobicity of viable N-protein species across the mutant spectrum. The polarity index (A) and hydrophobicity index (B) was calculated based on amino acid composition for all distinct sequences of N-FL, the folded domains (NTD and CTD), and the IDRs (N-arm, linker, and C-arm). Further subdivisions of the linker into the SR-rich and L-rich regions, and subdivisions of the C-arm into the N3 region and the C-terminal remainder of the C-arm (C-arm1) are indicated in the arrows. Highlighted by horizontal lines are the values for the corresponding peptides from the ancestral sequence Wuhan-Hu-1 (blue), and including the defining mutations of the Delta variant (dotted red) and the Omicron variant (dashed green), respectively. Symbols indicate values for SARS-CoV-2 (ancestral reference, light blue circles), and corresponding peptides from SARS-CoV-1 (red up triangles), MERS (red down triangles), MHV (red squares), human coronavirus NL63 (grey pentagrams), and the bat coronavirus APD51511.1 (grey diamonds).

It is useful to subdivide the linker IDR further to distinguish the SR-rich region (N:175-205), which exhibits high polarity and low hydrophobicity, from the L-rich region (N:206-247), which exhibits opposite behavior and is among the sequence stretches with lowest polarity values and highest hydrophobicity (**Figure 2**, red arrows in magenta shaded columns). Despite significant spread across the mutant spectrum, there is no overlap in these properties, which suggests biophysical constraints require the distinct polar and non-polar properties of the SR-rich region and the L-rich region, respectively. Indeed, these regions in the linker IDR have been recognized to play distinct functional roles: The SR-region provides a major hub for phosphorylation, aids in NA-binding, and mediates NA-binding induced allosteric interactions between NTD and the L-rich region (Pontoriero et al., 2022; Yaron et al., 2022; Zhao et al., 2023). This is distinct from the L-rich region, which has a propensity for the formation of transient helices that interact with NSP3 (Bessa et al., 2022), and can assemble *via* hydrophobic interactions to from coiled-coiled oligomers that contribute to the architecture of RNPs in viral assembly (Adly et al., 2023; Zhao et al., 2024, 2023).

Similarly, the C-arm IDR can be subdivided in the N3 region (N:390-419) and the remainder (‘C-arm1’, N:364-389), which also have strikingly different properties (**Figure 2**, blue arrows in cyan shaded columns): Whereas the connecting C-arm portion is by far the most polar, the N-terminal N3-region is among the most hydrophobic regions of the entire protein. Interestingly, the N3 region contains a transient helix (Cubuk et al., 2021; Zhao et al., 2023, 2022), which may be involved in recognition of the packaging signal and M-protein interactions localized here (Kuo et al., 2016; Masters, 2019). Again, the difference in the physicochemical properties of these regions persists throughout the entire ensemble of sequences despite their significant spread and high mutation frequencies (**Figure 1B**).

Charges in proteins can control multiple properties related to electrostatic interactions, from functions of active sites to protein solubility, protein interactions, and conformational ensembles in IDRs (Garcia-Viloca et al., 2004; Gerstein and Chothia, 1996; Gitlin et al., 2006; Mao et al., 2010). The net charges of the different N-protein regions are displayed in **Figure 3A**. Similar to polarity and hydrophobicity, viable sequences can have significant spread of net charges among all the mutants, amounting to departures by ±(1-2) from the ancestral sequence. This is expected considering the replacement and introduction of charged residues in the mutational landscape, for example, including those from the defining substitutions of variants. The positive charge of the overall basic protein is shared similarly among all folded domains and IDRs. However, noteworthy is again the contrast arising from subdivision of the linker and C-arm, which displays uneven and non-overlapping distributions: despite the strongly basic character of the linker, its L-rich sequence is nearly neutral; similarly, the basic C-arm splits into an even more basic C-arm1 and an acidic N3 tail region. These differences are highly significant and persist throughout the mutant spectrum.

**Figure 3.**
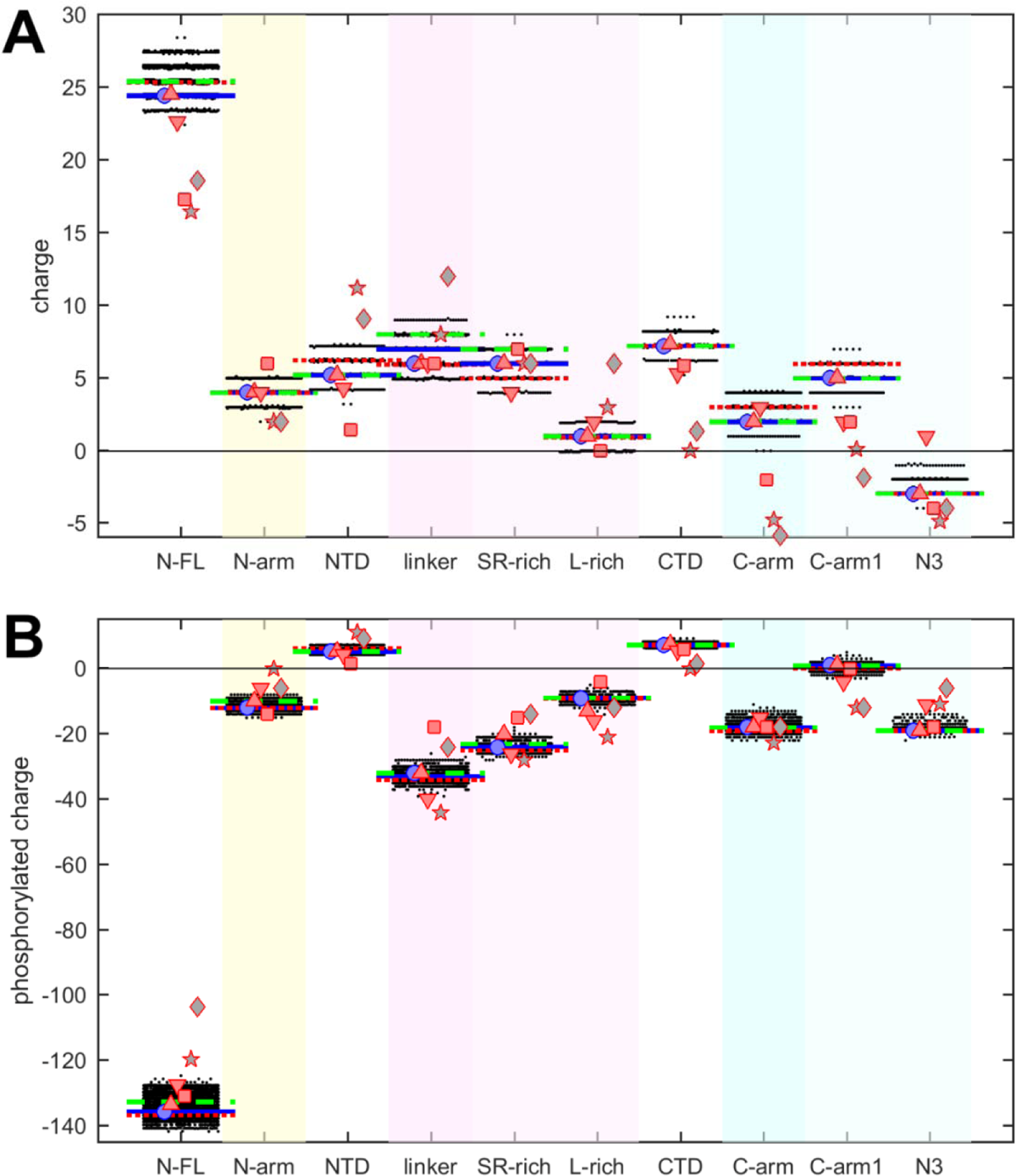
Beehive plots showing the distributions of charges of viable N-protein species. (A) Charges were calculated based on the amino acid composition of different N-protein regions as in Figure 2. Highlighted by horizontal lines are the values for the corresponding peptides from the ancestral sequence Wuhan-Hu-1 (blue), and including the defining mutations of the Delta variant (dotted red) and the Omicron variant (dashed green), respectively. Symbols indicate values for SARS-CoV-2 (ancestral sequence, blue circles), SARS-CoV-1 (red up triangles), MERS (red down triangles), MHV (red squares), NL63 (grey pentagrams), and bat coronavirus APD51511.1 (grey diamonds). (B) Same as in (A), with added charges from maximally phosphorylated serine, threonine, and tyrosine residues in the IDRs.

It is well-established that intracellular N-protein can be heavily phosphorylated (in contrast to N-protein in the virion) (Botova et al., 2024; Carlson et al., 2020; Fung and Liu, 2018; Johnson et al., 2022; Yaron et al., 2022). As reviewed in (Yaron et al., 2022), most serine, threonine and tyrosine residues in the disordered regions (30 of 37) have been found phosphorylated in different proteomic analyses. Accordingly, we estimated the maximum charge when all of these residues in the IDRs are phosphorylated (**Figure 3B**). This leads to a negative charge for all IDRs. As might be expected, the largest impact was found in the SR-rich region of the linker, which carries the highest density of phosphorylation sites. Interestingly, despite the considerable spread of net charges within families of mutant sequences, the differences between the regions remain highly significant.

It is noteworthy that the defining mutations of the Delta- and Omicron-variant (denoted by dotted red and dashed green horizontal lines, respectively) do impact the hydrophobicity, polarity, and charges in all of the N-protein regions. However, their values do not stand out from the clouds of values across the mutant spectrum, which include more extreme values throughout.

### Physicochemical properties of related coronaviruses

The distinct physicochemical properties of the linker and C-arm sub-segments persist throughout the mutant spectrum, which suggests these constitute biophysical constraints for functional SARS-CoV-2 N-protein. Therefore, we asked whether this holds true for N-protein from related coronaviruses such as SARS-CoV-1 (P59595.1), Middle East respiratory syndrome coronavirus (MERS, YP_009047211.1), murine hepatitis virus (MHV, NP_045302.1), human coronavirus NL63 (Q6Q1R8.1), and the 229E-related bat coronavirus APD51511.1. To this end, we used alignment of their consensus sequences to SARS-CoV-2 N-protein (shown previously (Zhao et al., 2022)) to subdivide all N-proteins into equivalent regions (**Supplemental File S1**). As shown in **Table 1**, the resulting peptides present high sequence identity scores for the FL protein and the folded domains, but, with exception of SARS-CoV-1, have little to no sequence identity in the IDRs. This observation is consistent with the high mutation frequency of the IDRs.

**Table 1.** Sequence alignment score of segments from related coronaviruses.

| virus | full-length | N-arm | NTD | linker | SR-rich | L-rich | CTD | C-arm | C-arm1 | N3 |
| --- | --- | --- | --- | --- | --- | --- | --- | --- | --- | --- |
| SARS-CoV-1 | 672<sup>(1)</sup> | 68.6 | 263 | 41.6 | 44.7 | 30 | 231 | 60.5 | 75.3 | 77 |
| MERS | 276 | 13.9 | 157 |  |  |  | 112 | 14.6 | 23.5 |  |
| MHV | 192 |  | 114 | 14.6 |  |  | 80.5 | 14.6 | 13.4 |  |
| NL63 | 67.4 |  | 58.9 |  |  |  | 61.6 |  |  |  |
| APD51511.1 | 61.2 |  | 44.3 |  |  |  | 44.3 |  |  |  |

<sup>(1)</sup> Values are BLASTp total alignment scores

The resulting peptides were subjected to the same analyses of physicochemical properties described above for SARS-CoV-2 N-protein. The results are displayed in **Figures 2 and 3** as symbols. With regard to hydrophobicity (**Figure 2B**), the FL proteins and folded domains show values within the range of the SARS-CoV-2 mutant spectrum. By contrast, more significant spread is observed in most IDR peptides. Nonetheless, the pattern observed for SARS-CoV-2 of hydrophobicity and polarity values of IDRs relative to those of the folded domains, and the pattern comparing subdivisions of the IDRs is closely mirrored for SARS-CoV-1, MERS, and MHV (red symbols). Similar patterns, although with some divergence, are observed for the NL63 and APD51511.1 IDRs (grey pentagrams and diamonds, respectively) which have the least sequence identity to SARS-CoV-2.

Polarity values (**Figure 2A**) of all coronavirus linker peptides are higher than either their corresponding FL, NTD, or CTD regions. The subdivision of the linker in the peptides corresponding to SR-rich and L-rich regions of SARS-CoV-2 follow the same qualitative trend, with higher polarity in the equivalent SR-rich and lower polarity in the equivalent L-rich peptides for all coronaviruses studied. Similarly, the properties of the equivalent C-arm and subdivision of C-arm1 and N3 peptides for SARS-CoV-1, MERS, and MHV (red symbols) closely track the values from the mutant spectrum of SARS-CoV-2, although this is not the case for the more distant NL63 and APD51511.1 (grey symbols).

Charge properties of related coronaviruses follow a similar pattern of SARS-CoV-2 (**Figure 3A**), although with somewhat greater differences, particularly again for NL63 and APD51511.1. Peptides corresponding to L-rich regions exhibit low charge, distinctly below those of the SR-rich regions, and similarly, N3 peptides have lower charges than C-arm-1 peptides of the corresponding viral species, and nearly all are acidic. Even though it is unclear to what extent IDRs of other coronaviruses can be phosphorylated, their amino acid composition would provide similar potential as SARS-CoV-2, as the completely phosphorylated charges of all peptides follow closely those of SARS-CoV-2 (**Figure 3B**).

This suggests that the charge properties and phosphorylation, like polarity and hydrophobicity, of the equivalent IDR sub-regions are functional biophysical constraints maintained across related coronaviruses despite little sequence conservation.

### Biophysical properties of select mutants

Unfortunately, it is impossible to express and experimentally characterize biophysical properties of all mutant species. Therefore, to assess the range of phenotype variation we examine only six exemplary protein constructs related to variants of concern in comparison with the Wuhan-Hu-1 reference molecule, N_ref_ (**Table 2**): 1) N:R203K/G204R with a double mutation in the disordered linker that arose early in the Alpha-variant (B.1.1.7), but occurs also in the Gamma-variant (P.1), and all Omicron-variants (BA.1 through BA.5). It was found to modulate phosphorylation of cytosolic N-protein, enhance assembly in a VLP assay, and increase viral fitness (Johnson et al., 2022; Syed et al., 2023, 2022); 2) N:P13L/Δ31-33 carrying the mutation P13L and the deletion Δ31-33 that are part of the defining mutations of all Omicron variants, with P13L epidemiologically ranked as the most statistically significant N-protein mutation linked to increased fitness (Obermeyer et al., 2022; Oulas et al., 2021); 3) N_o_ is a combination of N:R203K/G204R and N:P13L/Δ31-33, carrying thereby the complete set of defining mutations of the BA.1 Omicron-variant; 4) N:G215C with a key mutation in the disordered linker that was associated with the rise of the 21J clade of the Delta-variant, and found to modulate a transient helix in the L-rich linker region (Zhao et al., 2022); In a reverse genetics system, N:G215C was recently reported to cause significantly increased viral growth and altered virion morphology (Kubinski et al., 2024). 5) N:D63G containing another defining mutation of the Delta-variant, located in the NTD and epidemiologically ranked above G215C in increasing SARS-CoV-2 fitness (Obermeyer et al., 2022); and 6) N_δ_ carrying all four defining mutations D63G, R203M, G215C, D377Y of the Delta-variant. As detailed in **Table 2**, all of these species are found in the genomic database, and in combination with additional mutations occur in a high fraction of all genomes (exceeding the frequency of the ancestral Wuhan-Hu-1 N-protein by an order of magnitude). However, with the exception of N:G215C, none of the mutants has been studied in detail with regard to their macromolecular biophysical properties.

**Table 2.** Overview of N-protein species compared in biophysical experiments.

| designation | N-protein mutations | *n* exclusive instances<sup>(1)</sup> | occur in # of distinct sequences<sup>(2)</sup> | occurs in % of all genomes<sup>(3)</sup> | in set of defining VOC mutations<sup>(4)</sup> |
| --- | --- | --- | --- | --- | --- |
| N:R203K/G204R | R203K, G204R | 53,282 | 17,552 | 57% | α, γ, o |
| N:P13L/Δ31-33 | P13L, Δ31-33 | 9,548 | 12,503 | 47% | o |
| N<sub>o</sub> | P13L, Δ31-33, R203K, G204R | 791,613 | 10,238 | 46% | o (all BA.1)<sup>(5)</sup> |
| N<sub>δ</sub> | D63G, R203M, G215C, D377Y | > 1.2×10<sup>6</sup> | 9,397 | 33% | δ (all 21J)<sup>(5)</sup> |
| N:G215C | G215C | 60 | 10,562 | 34% | δ |
| N:D63G | D63G | 182 | 12,443 | 36% | δ |
| N<sub>ref</sub> | none | 38,929 | NA | 3.6% | NA |

<sup>(1)</sup> Number of genomes where the indicated mutations are the only N mutations. <sup>(2)</sup> Number of unique N-protein sequences in which indicated mutations are present, alongside other mutations. <sup>(3)</sup> Percentage of all sequenced genomes carrying the specific mutation. <sup>(4)</sup> Variants of concern for which indicated mutations are part (or all) of the defining set of N-mutations. <sup>(5)</sup> These sets of mutations comprise all defining N-protein mutations of this variant. <sup>(5)</sup> Literature on definition or biophysical characterization of the mutant.

All mutations considered here are within the IDRs, except for N:D63G, a mutation characteristic of the Delta variant. The presence of the N:D63G mutation in the NTD is highlighted in the shift of the intrinsic fluorescence quantum yield of this mutant in comparison to N_ref_ (**Figure 4A**). This may be attributed to changes in the local environment of tryptophan W108, which is partially surface exposed and structurally near the aspartic acid D63, as indicated by AlphaFold structural predictions (**Supplemental Figure S1**). D63G ablates a negative surface charge near the nucleic acid (NA) binding site of the NTD, which poses the question whether this mutation alters NA binding affinity. We assessed this using sedimentation velocity analytical ultracentrifugation (SV-AUC) with the oligonucleotide T_10_ as a NA probe. T_10_ is comparable in length to the NTD binding canyon for NA but does not permit multi-valent binding (Dinesh et al., 2020; Zhao et al., 2021). No significant differences in the intrinsic binding affinity to T_10_ was detected between N:D63G, other mutants, and the ancestral species (**Supplemental Figure S2**).

**Figure 4.**
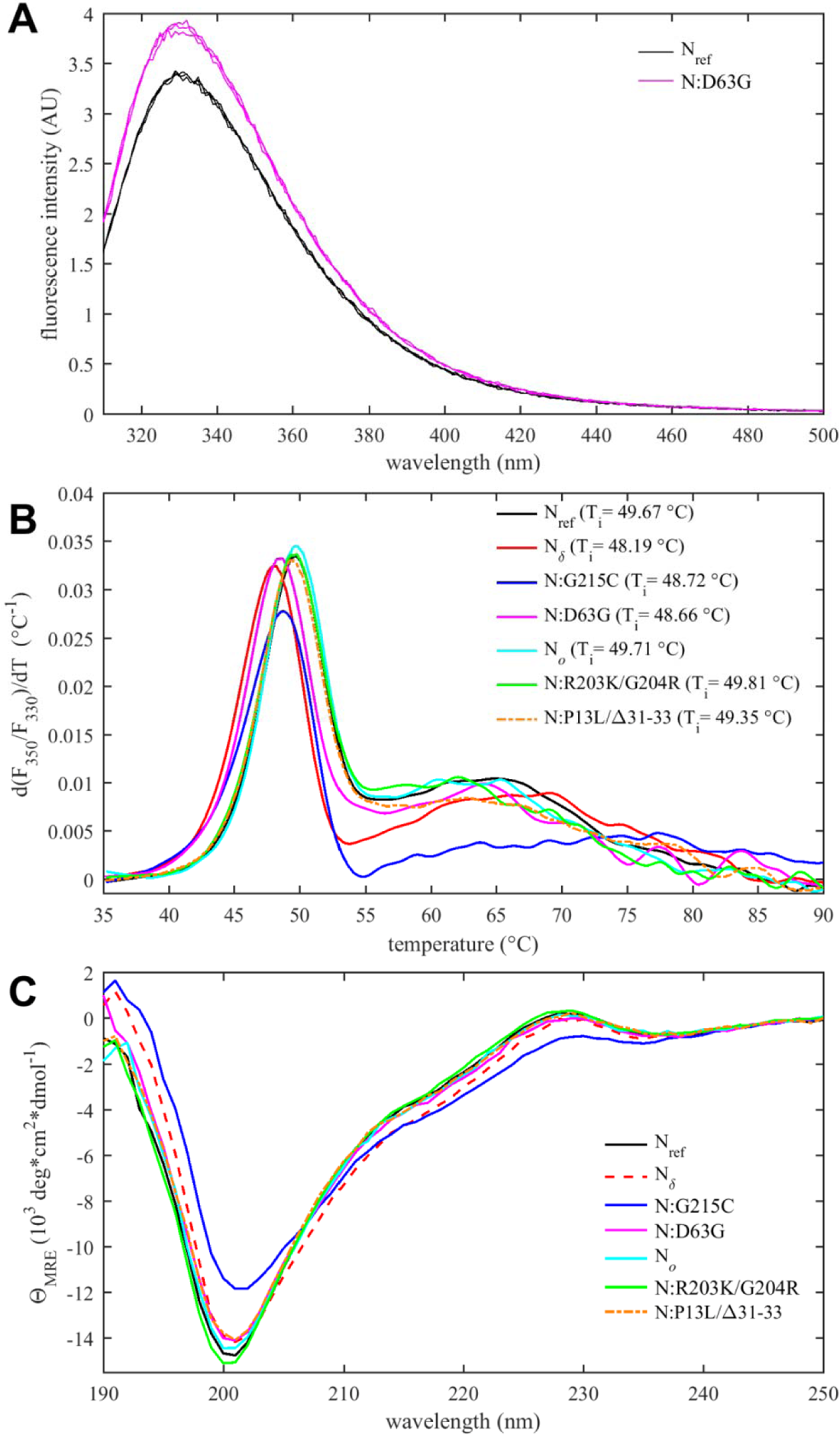
Thermodynamic stability and structural differences of N-protein reference and mutant species. (*A*) Intrinsic fluorescence spectrum of N:D63G in comparison with N_ref_, showing spectra in triplicate. (*B*) Differential scanning fluorometry, with the temperature of maximum fluorescence ratio derivative (*T*_i_-values, with an estimated precision 0.3 °C).(*C*) Circular dichroism spectra of all N-protein species (spectra with error bars are shown in **Supplemental Figure S3**).

A parameter of great interest from an evolutionary perspective is the thermal stability of the folded domains. This property can be assessed experimentally by differential scanning fluorometry (DSF), which reports on temperature-driven changes in the environment of aromatic amino acids due to changes in solvent exposure (Eftink, 2000). Such changes may occur during unfolding or as a result of other conformational changes. In the case of N-protein, conveniently all tryptophan and tyrosine residues of N-protein are located in the NTD and CTD, such that changes in the intrinsic fluorescence report exclusively on changes in the state of the folded domains. As shown in **Figure 4B**, a major transition is observed with an inflection point at *T*_i_ ≈ 49 °C. Compared to the reproducibility of transition temperatures of ±0.3 °C, significant shifts from the ancestral N-protein can be discerned: While Omicron mutations N_o_, N:R203K/G204R, and N:P13L/Δ31-33 are neutral, those occurring in the Delta variant (N:D63G, N:G215C, and N_δ_) are destabilizing, i.e., they lower the transition temperature. Interestingly, apparent destabilization of the folded domains occurs in N:G215C despite the absence of mutations in the folded domains – 215C being located in the middle of the linker IDR. This non-local mutation effect points to altered intra-molecular interactions between IDRs and the folded domains, and/or changes in contacts between folded domains mediated through an altered oligomeric state. (This is corroborated in non-natural point mutants N:L222P and N:L222P/R226P which abrogate linker helix oligomerization (Zhao et al., 2023) and exhibit *T*_i_-values of ≈51 °C.) Furthermore, **Figure 4B** shows additional transitions occur at higher temperatures broadly in the range of 60 – 70 °C. While their origin is unclear, this signal may accompany the formation of higher-order structure. It is noteworthy that N:G215C is also distinctly different in this feature.

Secondary structure information from the entire molecule including the IDRs can be extracted from circular dichroism (CD) spectra. As may be observed from **Figure 4C** (and in more detail **Supplemental Figure S3**), significant variation occurs both in the magnitude of the negative ellipticity at ≈200 nm, which mainly reflects disordered residues, as well as the magnitude of the negative ellipticity at ≈220 nm, which reports on helical structure. Compared to the ancestral N_ref_, significantly less disorder and greater helicity is observed for N:G215C (and to lesser extent also for N_δ_), whereas slightly more disorder is indicated for N:R203K/G204R. Little difference to the ancestral molecule is observed for N_o_, N:P13L/Δ31-33, and N:D63G. The absence of significant changes for N:D63G is consistent with this mutation having only a subtle, if any, impact on the NTD conformation. For N:G215C, increased helicity can be attributed to the stabilization of transient helices in the leucine-rich region of the central linker IDR, as shown previously (Zhao et al., 2023, 2022).

Tertiary and quaternary structure can be assessed by sedimentation velocity analytical ultracentrifugation (SV-AUC) (**Figure 5A**). As reported previously, the ancestral N-protein at micromolar concentrations in NA-free form is a tightly linked dimer sedimenting at ≈4 S, without significant populations of higher oligomers (Forsythe et al., 2021; Ribeiro-Filho et al., 2022; Tarczewska et al., 2021; Zhao et al., 2022, 2021). The same behavior is observed for N:D63G, N_o_, N:R203K/G204R, as well as N:P13L/Δ31-33 at low micromolar concentrations (**Figure 5A**). By contrast, the G215C mutation promotes the formation of higher oligomers via stabilization of coiled-coil interactions of transient helices in the L-rich linker region (Zhao et al., 2023, 2022). This is consistent with the enhanced helical content of this mutant (**Figure 4C**). Oligomerization beyond the dimeric N_ref_ is also observed for N_δ_, which incorporates the 215C mutation, but less than for N:G215C. This is consistent with the intermediate helical content of N_δ_ observed in CD. Of the three additional mutations of N_δ_ relative to N:G215C, we speculate that D63G does not impact dimerization (as in N:D63G, **Figure 5A**), and that therefore either the distant D377Y and/or R203M might cause this reduction of helicity and oligomerization relative to N:G215C, noting that R203M is proximal to the L-rich region (215-235) reshaped by 215C (Zhao et al., 2023).

**Figure 5.**
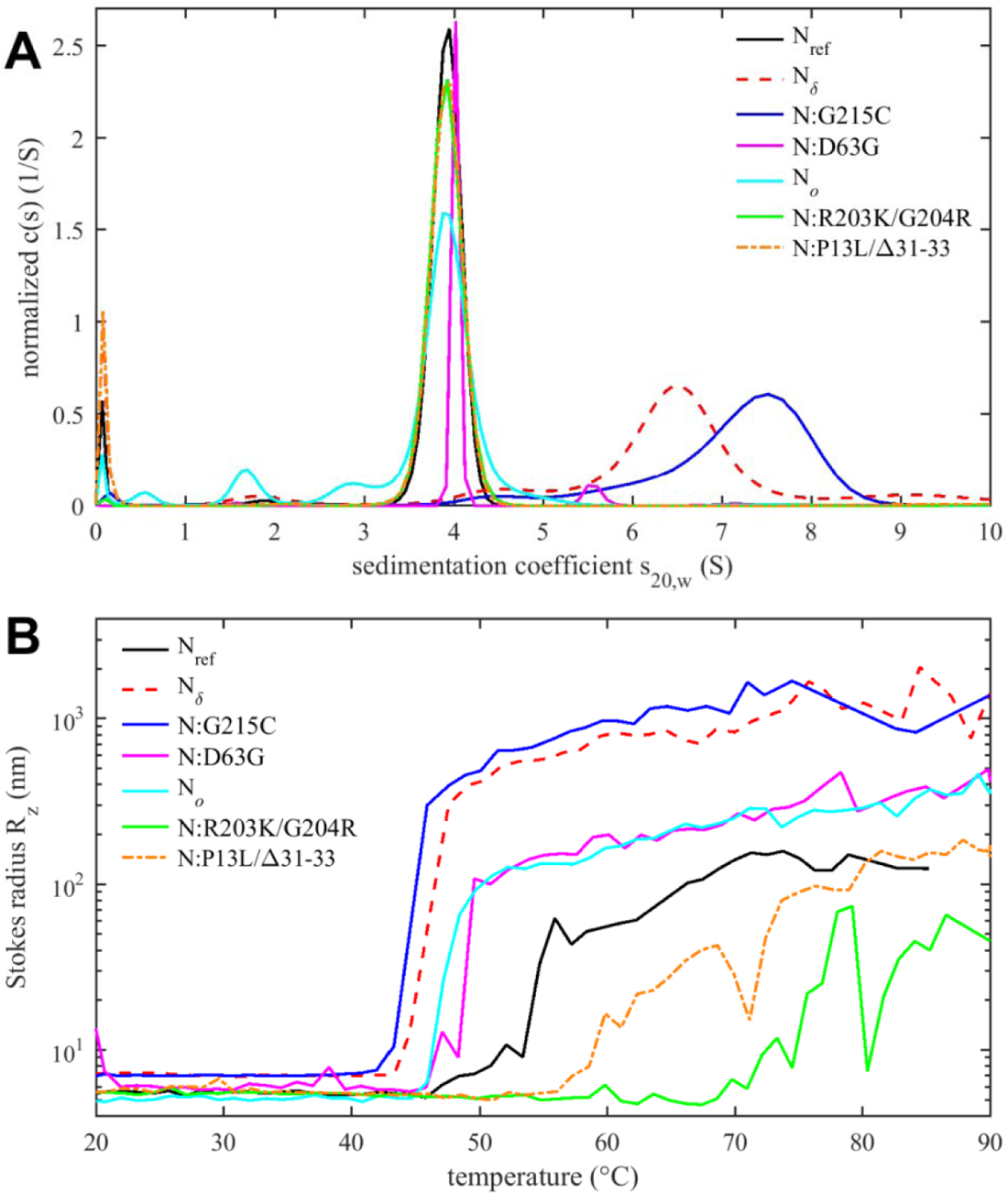
Tertiary and quaternary structure of N-protein species. (A) Sedimentation coefficient distributions *c*(*s*) from SV-AUC experiments show ≈4S dimers and higher oligomers. Data for N:G215C and N_δ_ are reproduced from (Zhao et al., 2022). (B) Temperature-dependent particle formation reported as average Stokes radius measured by dynamic light scattering.

As outlined in the introduction, N-protein has a propensity to form large particles and undergo LLPS (Carlson et al., 2020; Cascarina and Ross, 2022; Cubuk et al., 2021; Iserman et al., 2020; Jack et al., 2021; Lu et al., 2021; Perdikari et al., 2020; Savastano et al., 2020), which can be promoted at higher temperatures (Iserman et al., 2020; Zhao et al., 2021). **Figure 5B** shows the *z*-average particle size measured by dynamic light scattering (DLS) as a function of temperature. Particle formation is governed by a combination of processes, including the hydrophobicity-driven stabilization of the linker helix and its self-association, ultra-weak interactions across the entire protein contributing to LLPS, and unfolding and aggregation processes. This complicates a comparison of the temperature transitions observed in DSF (**Figure 4B**) and DLS (**Figure 5B**) (and a further technical difficulty may be potential differences in temporal lag of conformational rearrangements *versus* particle assembly kinetics).

Nevertheless, several clear observations can be made. As reported previously, N_ref_ forms clusters and particles at >55 °C (Zhao et al., 2021), which is strongly enhanced and occurs at a lower temperature for N:G215C, due to the enhancement of the linker oligomerization (**Figure 5B**) (Zhao et al., 2023). Very similar behavior is observed for N_δ_, which suggests that at higher temperatures any potential inhibitory role suspected of the R203M mutation on self-association may be less relevant compared to G215C. It is interesting to note that, correspondingly, both show a lower *T*_i_ in DSF. More moderate enhancement of particle formation is observed for N:D63G, which shows an onset already at ≈50 °C and larger particle averages than the ancestral protein. This also correlates with its significantly lower *T*_i_ in DSF. Thus, even subtle structural changes (as shown in **Supplemental Figure S1**) can impact the assembly behavior.

The opposite effect, strong inhibition of particle formation, is observed for the N:R203K/G204R double mutant. Here, particles form only at temperatures > 70 °C, as a mixture of smaller clusters with some very large aggregates that adventitiously enter the light path in DLS and cause fluctuations in the *z*-average Stokes radius. Interestingly, although N_o_comprises the R203K/G204R mutation, N_o_does not share this behavior but instead exhibits slightly enhanced particle formation relative to the ancestral N_ref_, comparable to N:D63G. This points to the role of additional mutations in N_o_, which besides R203K/G204R features the N-arm mutations P13L and Δ31-33. Interestingly, by themselves in N:P13L/Δ31-33 the particle formation is also suppressed relative to N_ref_, although less so than for N:R203K/G204R. This again points to non-additive effects, suggesting that the combination of N-arm and linker IDR mutations in N_o_alter the effect of either set of inhibitory mutations alone, to jointly promote particle formation of N_o_.

We were curious whether IDR mutations might alter particle formation through modulation of existing or introduction of new protein-protein interfaces. We focused on Omicron mutations as these are obligatory an all currently circulating strains, and specifically on N-arm mutations, which have recently been implicated in altered intramolecular interactions with NA-occupied NTD (Cubuk et al., 2023). Even though SV-AUC showed no indication of self-association of N:P13L/Δ31-33 at low micromolar concentrations, weak interactions with K_d_ > mM would not be detectable under these conditions yet could be highly relevant in the context of multi-valent complexes (Zhao et al., 2024). Following the roadmap used previously for the study of the weak self-association of the leucine-rich linker IDR (Zhao et al., 2023), we restricted the protein to the N-arm peptide such that it can be studied at much higher concentrations. To this end, we compared solution behavior of the N-arm constructs N_ref_:(1-43) with the Omicron N-arm N:P13L/Δ31-33(1-43), as well as the N-arm with individual mutation N:P13L(1-43) and deletion N:Δ31-33(1-43). Unexpectedly, solutions of N:P13L/Δ31-33(1-43) exhibited elevated viscosity after storage for several days at 4 °C in 20 mM HEPES, 150 mM NaCl, pH 7.5. Since this is a tell-tale sign of weak protein interactions, we carried out ColabFold structural predictions. Even though ColabFold is trained to predict folded structures, it has been found to be frequently successfully in predicting transient folds in IDRs (Alderson et al., 2023; Zhao et al., 2023). Indeed, it predicts that replacement of proline at position 13 by leucine allows for formation of parallel sheets symmetrically arranged in higher-order N-arm oligomers (**Supplemental Figure S4**). We proceeded to test oligomerization of the N-arm constructs experimentally in hydrodynamic studies. **Figure 6A** shows autocorrelation functions of all peptides. While the reference N-arm N_ref_:(1-43) and the construct carrying the Δ31-33 deletion behave as expected for non-interacting peptides of this size, the N-arm constructs carrying the P13L mutation (in particular, the Omicron N-arm N:P13L/Δ31-33(1-43)) exhibit very large correlation times. This may be indicative of either formation of large particles or the presence of weak interaction networks as in gels. Similarly, in SV-AUC (**Figure 6B**) the ancestral reference and the Δ31-33 deletion mutant sediment as expected for non-interacting N-arm peptides (Zhao et al., 2023), whereas rapidly sedimenting, anomalously shaped boundaries with ≈100-fold larger sedimentation coefficient were observed for the Omicron N-arm and the construct carrying solely the P13L mutation. This unequivocally demonstrates the introduction of new protein self-association interfaces from the P13L mutation. They are weak and not apparent in studies of the full-length protein N:P13L/Δ31-33 at low micromolar concentrations, but oligomers can be populated at the ≈100-fold higher achievable concentrations of the peptides, which mirrors the concentration range for in vitro observation of interactions of the leucine-rich linker helices (Zhao et al., 2023).

**Figure 6.**
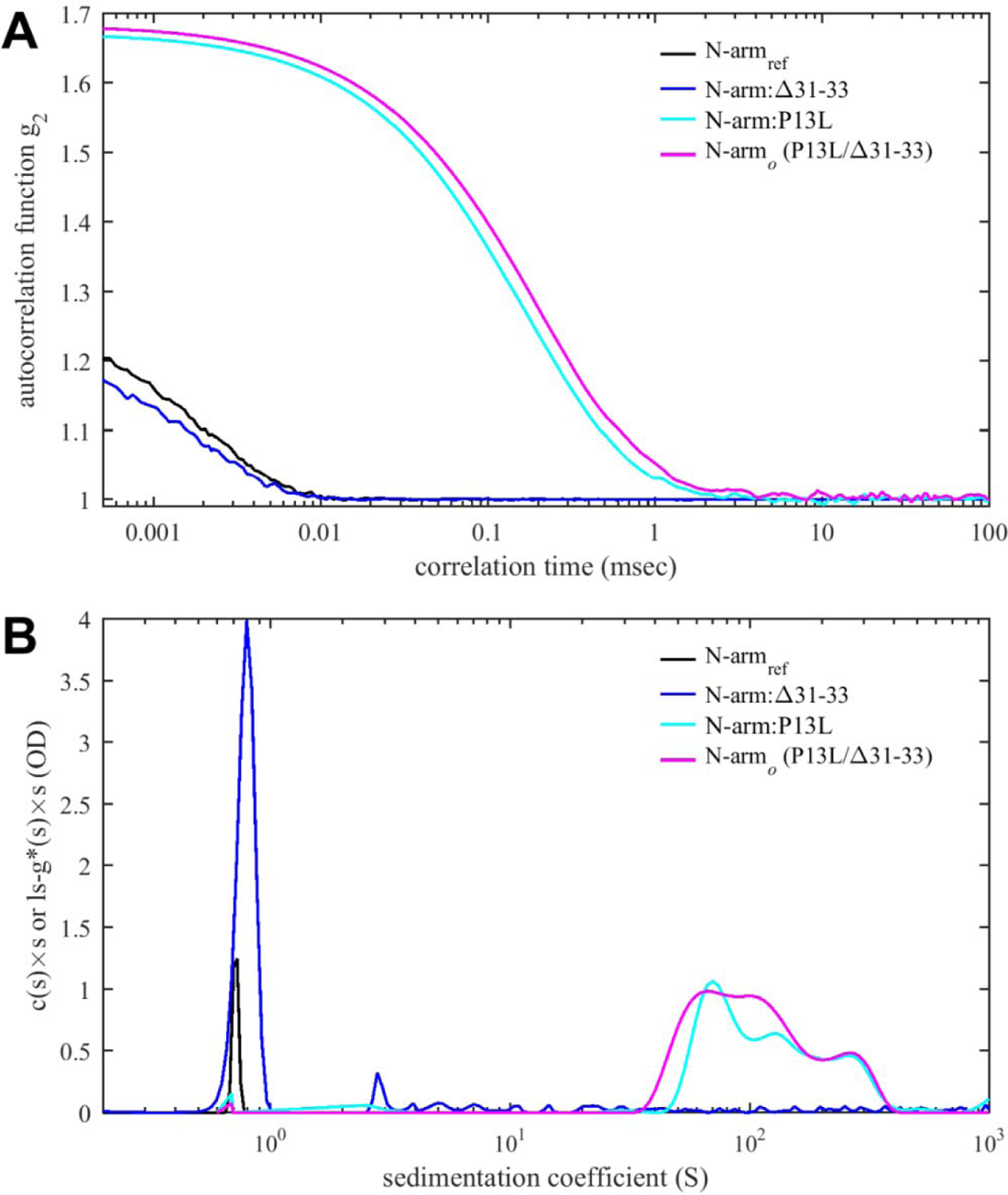
Protein-protein interactions of N-arm peptide containing the Omicron P13L mutation lead to large structures at high concentrations. (A) Autocorrelation functions from DLS (A) and sedimentation coefficient distributions from SV-AUC (B) for the ancestral reference N_ref_:(1-43) (black), N:Δ31-33(1-43) (blue), N:P13L(1-43) (cyan) and N:P13L/Δ31-33(1-43) (identical to the Omicron N-arm, magenta). All peptide concentrations are 400 µM, except for N_ref_:(1-43) in the SV-AUC experiment which is 275 µM, reproduced from previously reported data (Zhao et al., 2023).

The ability for N-protein to undergo LLPS in is thought to be crucial for several functions including interactions with stress granules, RNP assembly, and interactions with viral M-protein (Carlson et al., 2022; Cascarina and Ross, 2022; Iserman et al., 2020; Lu et al., 2021; Savastano et al., 2020). Weak protein-protein interactions and cluster formation such as shown in **Figure 5 and 6** can be coupled to LLPS, or alternatively LLPS may occur independent of clusters following Flory-Huggins theory (Kar et al., 2022). Therefore, we examined the impact of mutations on the propensity for LLPS. Images of phase separated condensates are shown in **Figure 7**, and corresponding histograms of droplet numbers and areas are shown in **Supplemental Figure S5**. As may be discerned from the top left panel of **Figure 7**, N_ref_ readily forms droplets in the presence of T_40_ oligonucleotides. Under the same conditions N:R203K/G204R (bottom left) does not display droplets, but forms few large particles with fibrillar morphology. In stark contrast, N:P13L/Δ31-33 (bottom center) readily forms droplets that appear to be more rapidly merging and growing than those of N_ref_ (**Supplemental Figure S6**). The combination of these mutations in N_o_ exhibits an intermediate propensity for LLPS with droplets in a dispersion of sizes. The most polydisperse distribution with largest droplets were observed for N:G215C (**Supplemental Figure S5**).

**Figure 7.**
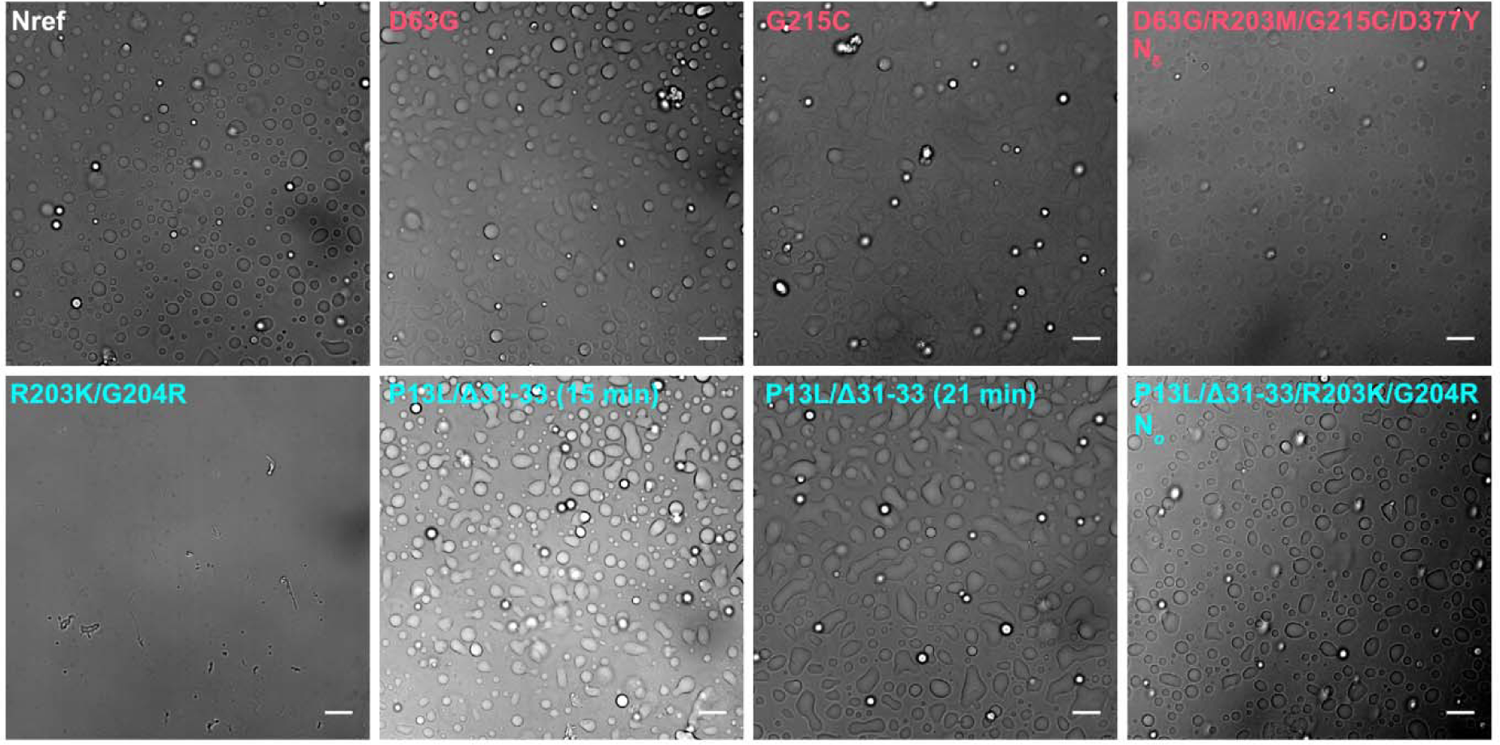
Differences in LLPS propensity of N-protein mutant species. Optical microscopy images were taken of 10 μM N-protein with 5 μM T_40_ (except N_δ_, which is 4 μM N-protein with 2 μM T_40_) in LS buffer after incubation for 15 min at room temperature. For N:P13L/Δ31-33 a second image was taken at the 21 min time point highlighting the growth of condensed phases. All scale bars are 10 µm. Histograms of particle areas are in **Supplemental Figure S5**, and a comparison of two time-points for Nref, N:R203K/G204R and N:P13L/Δ31-33 is provided in **Supplemental Figure S6**.

## Discussion

The SARS-CoV-2 pandemic has motivated the collection of virus genomic sequences on an unprecedented scale, which has generated invaluable data on the genomic diversity of an RNA virus. From the ensemble of observed consensus sequences of infected hosts we can extract, for the first time, an exhaustive map of possible amino acid replacements in viral proteins that are tolerable for viable virus (Bloom et al., 2023; Saldivar-Espinoza et al., 2023; Zhao et al., 2022). This brings into stark relief our limited understanding of the genotype/phenotype relationship, which is very detailed on some local functional aspects, such as spike protein antigenicity, but not much developed in general. This limits our ability to draw conclusions from the observed mutant spectrum on their variation in biophysical functions and fitness. Besides traditional sequence-based structure prediction and structure/function relationships, and more recent recognition of structural dynamics, new paradigms have emerged with increased understanding of the role of IDRs, their mimicry of short linear motifs, nonlocal physicochemical properties of sequence regions, and the ability of IDRs to promote macroscopic phase separation to generate or usurp condensates with virus-related functions. The extensive genomic data of SARS-CoV-2 presents an opportunity to probe how sequence diversity impacts these biophysical properties, and to examine what biophysical constraints exist for viral proteins to support viability. Focusing on SARS-CoV-2 N-protein we have studied the diversity of biophysical phenotypes with the goal to increase understanding of salient mechanisms of the many N-protein functions, and also to glean aspects of the biophysical fitness landscape underlying evolution.

On one hand, our studies of the diversity of nonlocal physicochemical properties of N-protein revealed the absence of tightly controlled hydrophobicity, polarity, and charges outside the folded domains. In the IDRs, individual mutations may alter each of these properties apparently without impacting viability, although modulatory fitness effects may be possible. For example, viable linker sequences span from 4.8 to 9.1 charges. On the other hand, a very clear separation of physicochemical parameters far exceeding mutational dispersion is maintained between the L-rich and SR-rich region of the linker IDR, and the N3 and remaining region of the C-arm IDR. These distinctions are likely functionally important, with the polarity and charges of the SR-rich linker region aiding in nucleic acid binding (Pontoriero et al., 2022), the hydrophobicity of the L-rich region aiding in assembly functions (Bessa et al., 2022; Zhao et al., 2024, 2023), and the acidic N3-region playing a role in NA- and M-protein interactions as suggested from analogy to MHV- and SARS-CoV-1 (Masters, 2019). These nonlocal features are also maintained in analogous consensus sequence regions of related coronaviruses, and thus provide further examples for nonlocal biophysical properties that are evolutionary conserved despite amino acid sequence divergence (Zarin et al. 2017; Zarin et al. 2021). It may seem as a paradox that despite this conservation these features seem not very fine-tuned and that significant variation of these properties is still observed within the viable mutant spectrum, for polarity and hydrophobicity significantly exceeding the spread of parameter values of the folded domains. However, as mentioned above, the differences between IDR regions that appear associated with biophysical functions are of significantly larger magnitude. The tolerance for the remaining comparatively smaller fluctuations in physicochemical parameters may be important to allow sufficient local variation in sequence space for additional functions to evolve, such as the emergence of SLiMs to manipulate the host/virus interface (Davey et al., 2011; Schuck and Zhao, 2023). Correspondingly, in a recent study of SLiMs variation across the mutant spectrum, we found the total number and detailed location of phosphorylation SLiMs to vary considerably in the SR-rich region, but to be maintained overall at a high level across this region (Schuck and Zhao, 2023).

Other nonlocal properties were studied experimentally, though unavoidably only by example of several different SARS-CoV-2 N-protein species. We selected conspicuous mutations in variants of concern, but each of the constructs studied also represents in itself viable N-protein species occurring in consensus sequences of the genomic database. Strikingly, point mutations can affect protein properties on all levels of organization, from thermodynamic stability and secondary structure to intra- and inter-molecular interactions, oligomeric state, particle formation, and LLPS. These results must be considered in the context of the highly dynamic nature of N-protein, which is caused by the flexibility of intrinsically disordered domains (Cubuk et al., 2023, 2021; Redzic et al., 2021; Zhao et al., 2021), the NTD and its disordered β-hairpin (Redzic et al., 2021), and the large-scale conformational fluctuations of the N-protein dimer in solution (Botova et al., 2024; Ribeiro-Filho et al., 2022; Różycki and Boura, 2022). High sequence plasticity is accompanied by high plasticity of protein configuration and delicate balances of protein interactions that can be significantly shifted by single mutations with nonlocal effects.

Our results highlight two different mechanisms through which mutation effects may be propagated across the protein. First, mutations can impact the transient helix in the hydrophobic L-rich region of the linker, and, as we have shown previously, promote its helical conformation and self-association into higher oligomeric states (Zhao et al., 2023, 2022). This, in turn, may impact collision frequency or other intra-molecular interactions with folded domains, such as the recently reported intra-molecular contact of the L-rich region to the NTD observed by NMR (Botova et al., 2024). This is reflected in the altered secondary structure observed in CD of N_δ_ and N:G215C, their oligomers observed in SV-AUC, and this would explain the impact of the G215C mutation on the thermal stability reported by intrinsic fluorescence localized to the NTD and CTD. In addition, changes near the L-rich transient helix also impact weak protein interactions and amplify to enhanced particle formation and altered LLPS. Notably, introduction of N:G215C in a reverse genetics system resulted in enhanced viral replication and larger virions (Kubinski et al., 2024).

Second, mutation frequencies peak in the downstream end of the SR-rich linker region, including the double mutation R203K/G204R that is part of the defining mutations of Omicron (and other) variants. In different VLP and cellular assays (Johnson et al., 2022; Syed et al., 2022), it has been shown to modulate N-protein phosphorylation and thereby the balance between replication and assembly, with contributions from an emerging alternate, truncated N-protein (210-419) that itself supports assembly (Adly et al., 2023; Leary et al., 2021; Mears et al., 2022; Syed et al., 2023). In the present study, we found that full-length N:R203K/G204R strongly opposes both temperature-driven particle formation and LLPS with oligonucleotides. Interestingly, this effect can be compensated for by the additional N-arm mutation P13L that is present in all Omicron variants. P13L itself has been identified epidemiologically as a the most important driver of fitness in N-protein (Obermeyer et al., 2022; Oulas et al., 2021), but its biophysical effects have not been previously studied. We identified a distinct self-association propensity of N-arm peptides carrying the P13L mutation, and enhanced LLPS propensity of full-length N-protein carrying the complete set of N-arm mutations in Omicron, N:P13L/Δ31-33. This is consistent with the partial ‘rescue’ of particle formation and full restoration of LLPS propensity we have observed in the N_o_molecule with the complete set of P13L/Δ31-33/R203K/G204R mutations defining N-protein from the BA.1 (B.1.1.529) Omicron-variant. It is interesting to note that the R203K/G204R mutation, the P13L mutation, and the P13L/Δ31-33 combination each can occur independently of each other in viable virus species, with 261 genomes in the database carrying only the P13L mutation, 9,548 only the combination P13L/Δ31-33, and >50,000 genomes exclusively the double mutation R203K/G204R, even though their more frequent coexistence (by approximately tenfold, in all of Omicron variants) might suggests epistatic interactions and a fitness advantage. Related, it was shown that the P13L mutation causes complete loss of recognition of a CD8+ T-cell epitope, which may cause T-cell evasion (de Silva et al., 2021), and provide an additional fitness effects of this mutation. Compensating effects between linker IDR and N-arm mutations highlight the nonlocal consequences of IDR mutations. They also highlight the difficulty of assigning variant properties and fitness effects to a single mutation, given the entangled effects among the sets of multiple mutations defining the variants of concern.

In summary, the importance of IDRs in viral evolution was recognized previously for several reasons. Their inherent flexibility makes them more permissible for amino acid changes, which is born out in the mutational landscape of SARS-CoV-2. As mentioned above, this makes them well suited for host adaptation through remodeling of host protein interaction networks, which is exemplified in the clusters of host-specific mutations located in IDRs of Dengue virus proteins (Charon et al., 2018; Dolan et al., 2021). Mimicry of eukaryotic SLiMs is ubiquitous (Davey et al., 2011; Hagai et al., 2014; Mihalič et al., 2023), and as we have shown recently, the sequence space of SARS-CoV-2 N-protein IDRs allows presentation of a large fraction of known eukaryotic SLiMs (Schuck and Zhao, 2023). In addition, nonlocal sequence-distributed physicochemical features of IDRs such as their charge and hydrophobicity have been demonstrated recently to mediate biological functions and present evolutionary constraints (Moses et al., 2023; Zarin et al., 2021). This principle also holds true in the distinct properties of linker and C-arm regions of SARS-CoV-2 N-protein. A related nonlocal physicochemical property of IDRs is their propensity for supporting LLPS (Abyzov et al., 2022; Brocca et al., 2020; Pappu et al., 2023), which plays a key role in different N-protein functions (Carlson et al., 2020; Cascarina and Ross, 2022; Roden et al., 2022; Savastano et al., 2020). Finally, here we have observed the ability of mutations in IDRs to modulate overall biophysical properties such as thermal stability, oligomeric state, and assembly properties. In SARS-CoV-2 N-protein IDRs, the latter are mediated via weak interactions in transiently folded structures. In addition, the high flexibility of the IDRs and their resulting high intra-chain contact frequencies (Botova et al., 2024; Różycki and Boura, 2022) may magnify non-local consequences of mutations. This endows viral protein IDRs with yet another level of variation of the biophysical phenotype that can impact evolutionary fitness. Exploiting the emerging mutational landscape and sequence space presents both a challenge and opportunity to explore the biophysical phenotype spectrum and thereby uncovers the salient functional principles of RNA-virus proteins.

## Materials and Methods

### Mutational landscape, sequence alignment, and prediction of physicochemical properties

The Wuhan-Hu-1 isolate (GenBank QHD43423) (Wu et al., 2020) was used as the ancestral reference. Sequence data were based on consensus sequences of SARS-CoV-2 isolates submitted to the GISAID as previously described (Schuck and Zhao, 2023; Zhao et al., 2022). Briefly, sequence data were downloaded on January 20, 2023 from Nextstrain (Hadfield et al., 2018) and 5.06 million high quality preprocessed sequences were included in the analysis. 746 sequences exhibiting insertions in the N-protein were omitted, as well as those with more than 10 deletions in N-protein and those represented in fewer than 10 genome instances.

The resulting sequence database was parsed for different unique sequences for N-proteins and different segments, using MATLAB (MathWorks, Natick, MA). Sequence hydrophobicity was calculated in RStudio (https://posit.co/) using the package PEPTIDES (Osorio et al., 2015) and polarity and charge using the package ALAKAZAM (Gupta et al., 2015). For maximally phosphorylated charge, −2 was added to the total charge for each serine, threonine, and tyrosine in the IDRs.

Alignment of SARS and related coronavirus sequences (SARS-CoV-1 P59595.1, MERS YP_009047211.1, MHV NP_045302.1, human coronavirus NL63 Q6Q1R8.1, and 229E-related bat coronavirus APD51511.1) was carried out with COBALT at NLM (Papadopoulos and Agarwala, 2007), as shown in (Zhao et al., 2022). This alignment was used to dissect related viruses into regions corresponding to the SARS-CoV-2 regions (N-arm, NTD, linker, SR-rich, L-rich, CTD, Carm, Carm1, N3) (**Supplemental File S1**). The resulting segments of the related viruses were subjected to analysis of physicochemical properties as described above. Sequence similarity of the corresponding regions relative to the SARS-CoV-2 regions was calculated using BLAST blastp suite (Altschul et al., 1997), using an expectation threshold of 0.9, word size 2, and BLOSUM63 scoring matrix.

### Structure prediction

Structural predictions for NTD and N-arm were carried out using ColabFold (Mirdita et al., 2022) and graphics were generated using ChimeraX (Pettersen et al., 2021).

### Proteins, peptides, and oligonucleotides

N:D63G and N:G215C were purchased from EXONBIO (catalog# 19CoV-N170 and 19CoV-N180, San Diego, CA), while N_ref_, N:R203K/G204R, N:P13L/Δ31-33, N_o_, and N_δ_ were expressed in house as described previously (Zhao et al., 2023, 2022). Briefly, the full-length protein with an N-terminal Tobacco Etch Virus (TEV) cleavage site and 6His tag was cloned into the pET-29a(+) expression vector and transformed into One Shot BL21(DE3)pLysS E. coli (Thermo Fisher Scientific, Carlsbad, CA). After cell lysis, the protein was bound to a Ni-NTA column, and unfolded and refolded to remove residual protein-bound bacterial nucleic acid (Carlson et al., 2020). After elution the 6xHis tag was cleaved and the protein purified by size exclusion chromatography. Greater than 95% purity of the proteins was confirmed by SDS-PAGE, and the ratio of absorbance at 260 nm and 280 nm of ∼0.50-0.55 confirmed absence of nucleic acid. The latter is important to eliminate higher order N-protein oligomers induced by nucleic acid binding (Carlson et al., 2020; Tarczewska et al., 2021; Zhao et al., 2021). For a subset of mutants, the protein sequence and mass were tested and confirmed by LC-MS/MS and LC-MS mass spectrometry, respectively. Biophysical experiments were preceded by dialysis in either high-salt buffer (HS) consisting of 20mM HEPES, 150mM NaCl, pH 7.5, or low-salt buffer (LS) consisting of 10.1 mM Na_2_PO_4_, 1.8 mM KH_2_PO_4_, 2.7 mM KCl, 10 mM NaCl, pH 7.4 as indicated below.

The oligonucleotide T_40_ was purchased from Integrated DNA Technologies (Skokie, IL), as purified by HPLC and lyophilized. N-arm peptides were purchased from ABI Scientific (Sterling, VA), as purified by HPLC, examined by MALDI for purity and identity, and lyophilized.

### Spectroscopy

CD spectra were acquired in a Chirascan Q100 (Applied Photophysics, U.K.), using cuvettes of 1 mm pathlength, and data acquisition with 1 nm steps and 1 sec integration time. Results are averages of 3 acquisitions, corrected for buffer background. Protein concentration was 3 µM in buffer LS, except N_o_in buffer HS.

For the acquisition of fluorescence spectra, protein samples at 1 µM were loaded into a quartz cuvette with 1.0 cm optical pathlength. Steady-state tryptophan (Trp) fluorescence emission spectra in the range from 305 nm to 500 nm were recorded in a spectrofluorimeter (QuantaMaster, Photon Technology) with excitation at 295 nm using a 1.0 nm increment. Scans were acquired in triplicate.

DSF was carried out in a Tycho instrument (Nanotemper, Germany) as previously described (Zhao et al., 2021). Briefly, 10 µL samples were aspirated in capillaries (TY-C001, Nanotemper, Germany), and intrinsic fluorescence was measured at 350 nm and 330 nm while the temperature was ramped from 35°C to 95 °C at a rate of 30 °C/min. The first derivative of the intensity ratio was calculated as a function of temperature. DSF experiments were carried out at protein concentrations of 2 µM in buffer LS, except for N:R203K/G204R which was measured in buffer HS. As a buffer control, the difference in *T*_i_ for N_ref_ in LS and HS buffer was measured and found to be within error of data acquisition (**Supplemental Figure S7A**).

### Hydrodynamic techniques

SV-AUC experiments were carried out in a ProteomeLab XL-I analytical ultracentrifuge (Beckman Coulter, Indianapolis, IN) in standard configurations (Schuck et al., 2015), with instruments subjected to routine calibrations (Ghirlando et al., 2013). Briefly, 2 µM protein samples were filled in cell assemblies composed of charcoal-filled Epon double-sector centerpieces with sapphire windows, inserted in an 8-hole AN-50 TI rotor and temperature equilibrated. After acceleration to 50,000 rpm data acquisition commenced using the absorbance optical detector at 280 nm and the interference optical detector. Data were analyzed in SEDFIT (sedfitsedphat.nibib.nih.gov/software) in terms of a sedimentation coefficient distribution *c*(*s*) (Schuck, 2016). Proteins for self-association studies were in buffer HS, except N_ref_, N_δ_, and N:G215C were in LS, the latter causing a ≈5% increase in *s*-value (**Supplemental Figure S7B**). Typical accuracy of *c*(*s*) peaks are on the order of ≈1% for peak *s*-values and ≈1-2% for relative peak areas (Zhao et al., 2015).

Nucleic acid binding experiments were analyzed in buffer HS and LS with isotherms of signal weighted-average sedimentation coefficients in SEDPHAT (Schuck and Zhao, 2017). For studies of the N-arm peptide species, 400 µM peptide samples were studied by gravitational sweep sedimentation using rotor speed steps of 3,000 rpm, 10,000 rpm, 40,000 rpm, and 55,000 rpm (Ma et al., 2016) and analyzed with a model for apparent sedimentation coefficient distributions *ls*-*g*\*(*s*) (Schuck, 2016) as a qualitative representation of rapidly migrating boundaries of N:P13L(1:43) and N:P13L/Δ31-33(1:43), or with *c*(*s*) distributions for N_ref_:(1:43) and N:Δ31-33(1:43).

Temperature-dependent DLS autocorrelation data of N-protein species were collected in a NanoStar instrument (Wyatt Technology, Santa Barbara, CA) equipped with a 658 nm laser and using a detection angle of 90°. 100 µL samples at 3 µL N-protein in LS buffer were inserted into a 1 µL quartz cuvette (WNQC01-00, Wyatt Instruments), with excess sample to prevent evaporation in the observation chamber. A temperature ramp rate of 1°C/min was applied with 5 sec data acquisitions and averaging 3 replicates for each temperature point. Data were collected and processed with the software Dynamics 7.4 (Wyatt Instruments) to determine the average hydrodynamic radius by cumulant analysis.

DLS studies of N-arm peptides were carried out in a Prometheus Panta (Nanotemper, Germany) instrument at 20°C. The samples were loaded into a capillary (Nanotemper PR-AC002) and ACFs were acquired using the 405 nm laser at the detection angle of 140°.

### Optical microscopy

Optical imaging of *in vitro* phase-separated condensates was carried out as described previously (Zhao et al., 2021). Briefly, reaction mixtures of N protein and T_40_ in buffer LS were combined and mixed immediately prior to imaging. 20 µL samples were transferred onto a glass-bottom 35 mm dish (catalog # Part No: P35G-1.5-20-C, MatTek) for imaging at room temperature. Images were acquired on a Nikon Ti-E microscope equipped with a 100X 1.49 NA oil objective lens (LIDA light engine, Lumencor, Beaverton, OR) and recorded with a Prime 95B camera (Teledyne Photometrics) with a pixel size of 110 nanometers. Images were background subtracted and contrast enhanced using MATLAB (Mathworks, Natick, MA).

The segmentation of different shapes in the brightfield images was performed with deep learning methods. Specifically, a pre-trained model (versatile) from StarDist Napari Plugin (Schmidt et al., 2018) was employed to segment the shapes with the following parameters: Input image scaling: 0.5, probability threshold: 0.2, overlap threshold: 0.2. The labels were imported into Fiji and LABKIT (Arzt et al., 2022) for manual verification and correction. For each segmented object, the area was measured in MATLAB.

## Acknowledgements

We thank Dr. Yan Li for carrying out mass spectroscopy experiments. This work was supported by the Intramural Research Programs of the National Institute of Biomedical Imaging and Bioengineering (ZIA EB000099-02) and the National Heart, Lung, and Blood Institute, National Institutes of Health. This work utilized the computational resources of the NIH HPC Biowulf cluster for sequence analyses.

