## Supplementary Figure S3 for "Modulation of Biophysical Properties of Nucleocapsid Protein in the Mutant Spectrum of SARS-CoV-2"

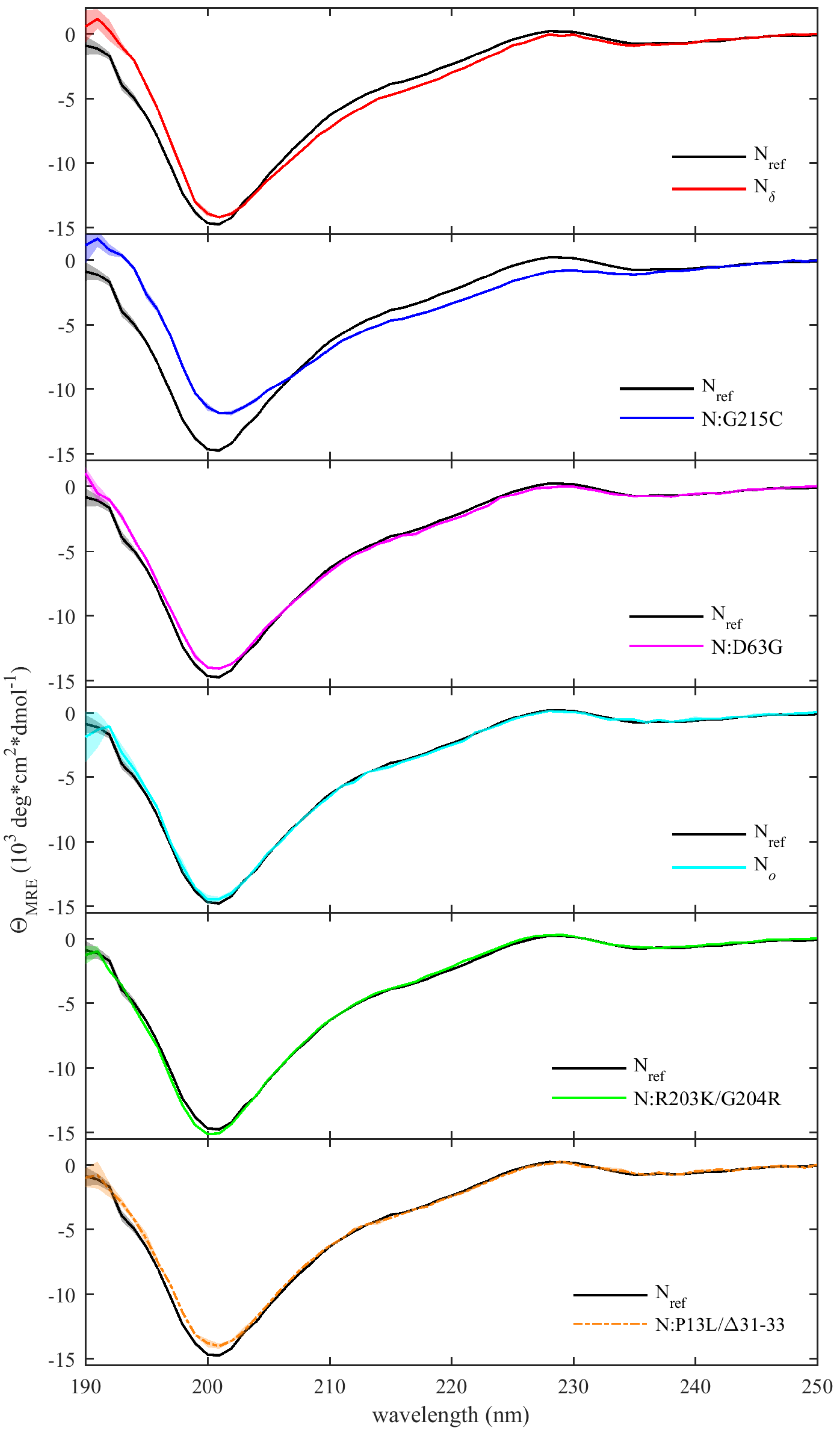


**Figure S3. Individual comparison of CD spectra.** The data from **Figure 4C** are reproduced and plotted in comparison with $N_{ref}$. Standard deviations from three acquired spectra are depicted as shaded bands.
