## Supplementary Figure S4 for "Modulation of Biophysical Properties of Nucleocapsid Protein in the Mutant Spectrum of SARS-CoV-2"

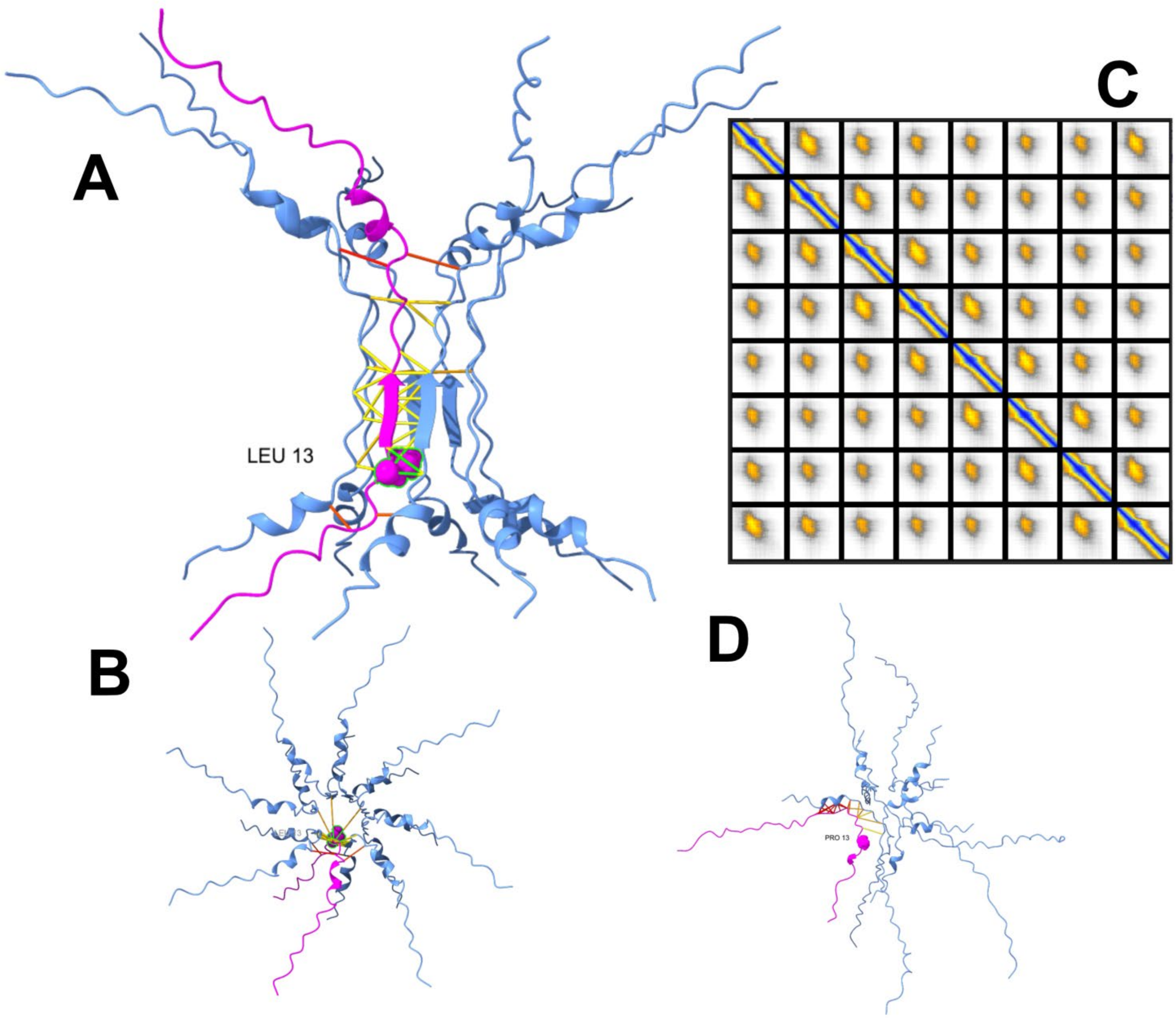


**Figure S4. Structural prediction of Omicron N-arm self-interactions.** (A) Best ColabFold prediction of eight Omicron N-arm (1:41) peptides with P13L and Δ31-33 mutations. For one chain shown in magenta, atoms of 13L are depicted and labeled, and contacts of this chain within 3.5 Å are color-coded by confidence. (B) Top view of (A). (C) Predicted alignment error (PAE) map showing symmetry and confidence of predicted interactions. (D) Best analogous prediction of ancestral N-arm interactions, highlighting the absence of order.
