## Supplemental Figure S5 for "Modulation of Biophysical Properties of Nucleocapsid Protein in the Mutant Spectrum of SARS-CoV-2"

**Supplementary Figure S5:**

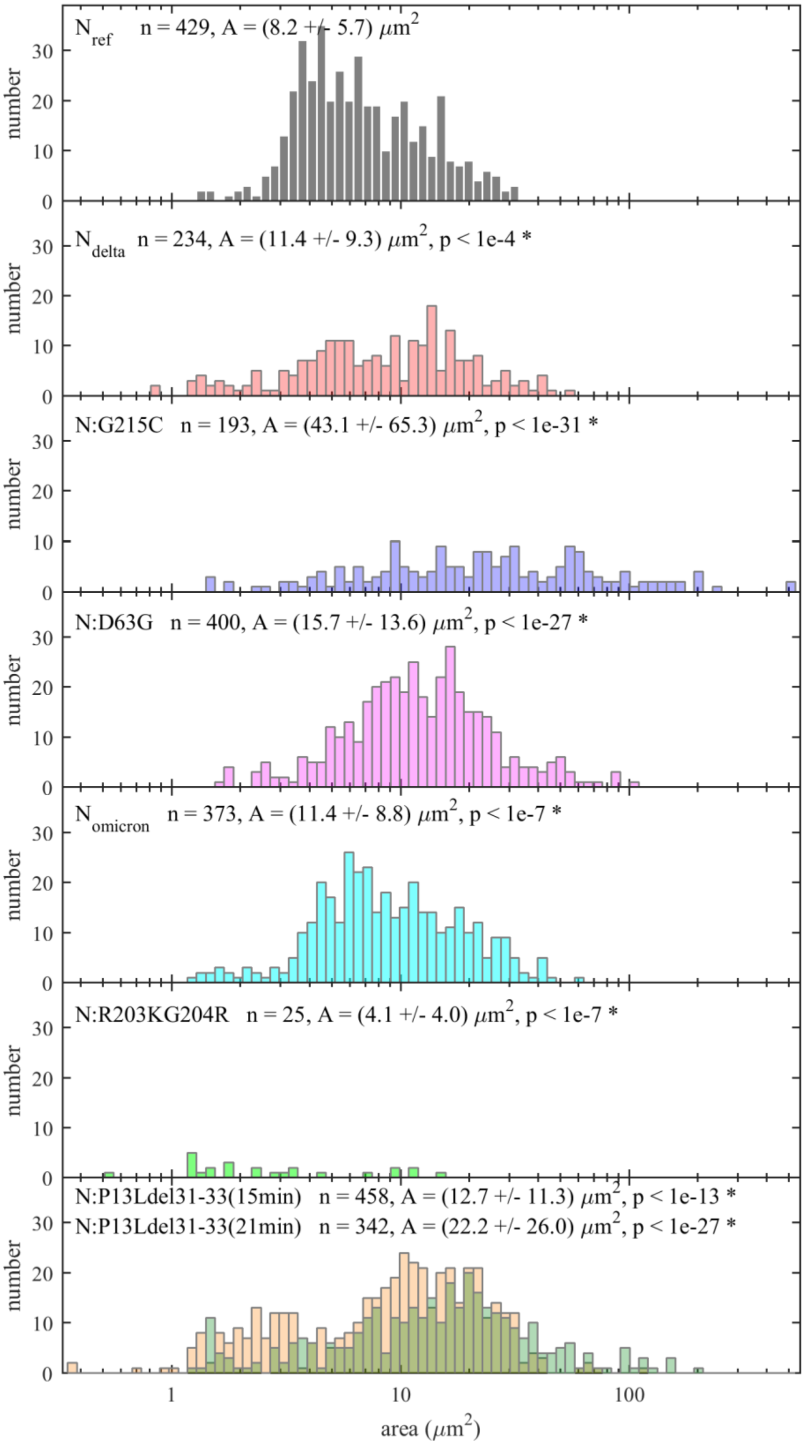


**Figure S5. Comparison of area distributions of droplets in images of Figure 7.** For each N-protein species, images were segmented to identify droplets. The values indicated are particle numbers, the mean area, the standard deviation of the area, and the probability that the sample is from the same distribution as $N_{ref}$ based on the two-sample Kolmogorov-Smirnov test.
