## Supplemental Figure S6 for "Modulation of Biophysical Properties of Nucleocapsid Protein in the Mutant Spectrum of SARS-CoV-2"

**Supplementary Figure S6:**

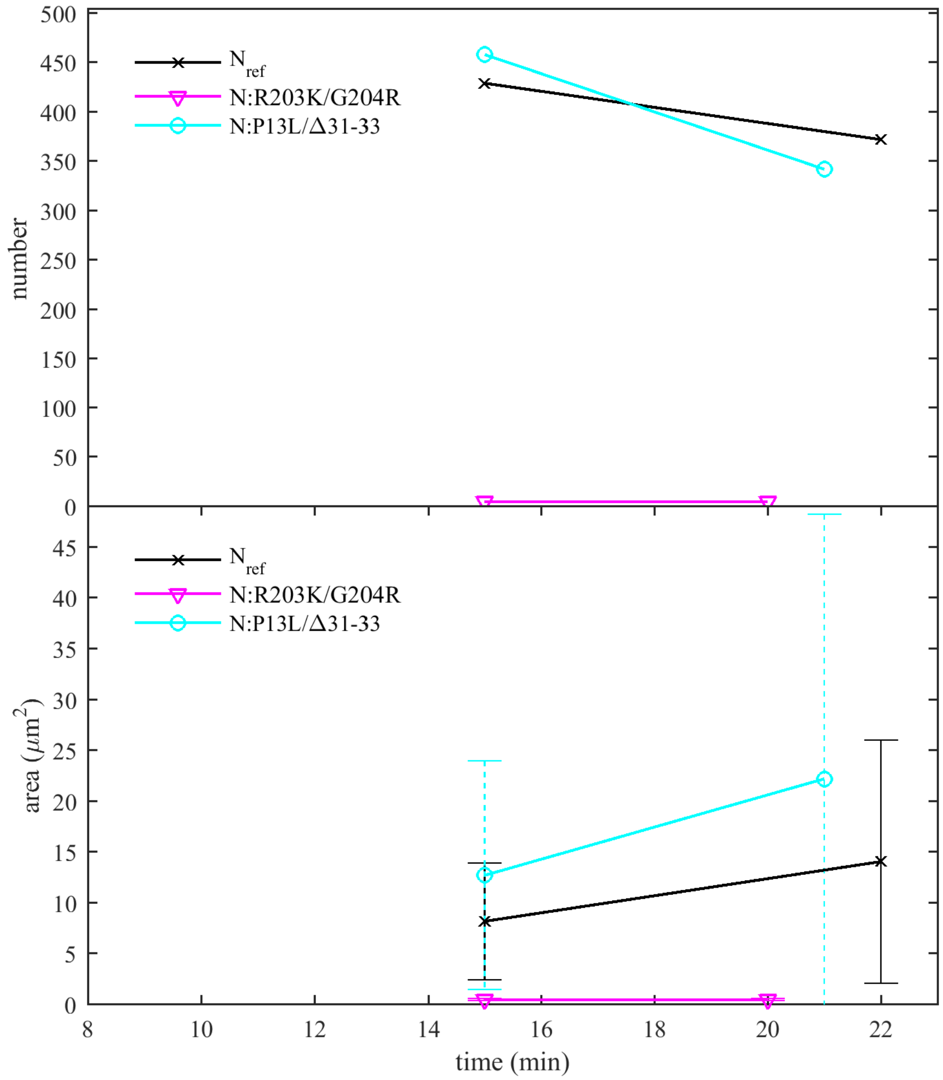


**Figure S6. Comparison of droplet area after LLPS at two points in time.** Similar to **Figure 7**, images of LLPS were recorded for $N_{ref}$, N:R203K/G204R, and N:P13L/Δ31-33 at two time-points for the same sample. The upper plot shows droplet numbers. The lower plot shows mean and standard deviations of the droplet area. Images and histograms for the early time points and the later time point of N:P13L/Δ31-33 are shown in **Figure 7** and **Supplementary Figure S4**, respectively.
