## Supplementary Figure S7 for "Modulation of Biophysical Properties of Nucleocapsid Protein in the Mutant Spectrum of SARS-CoV-2"

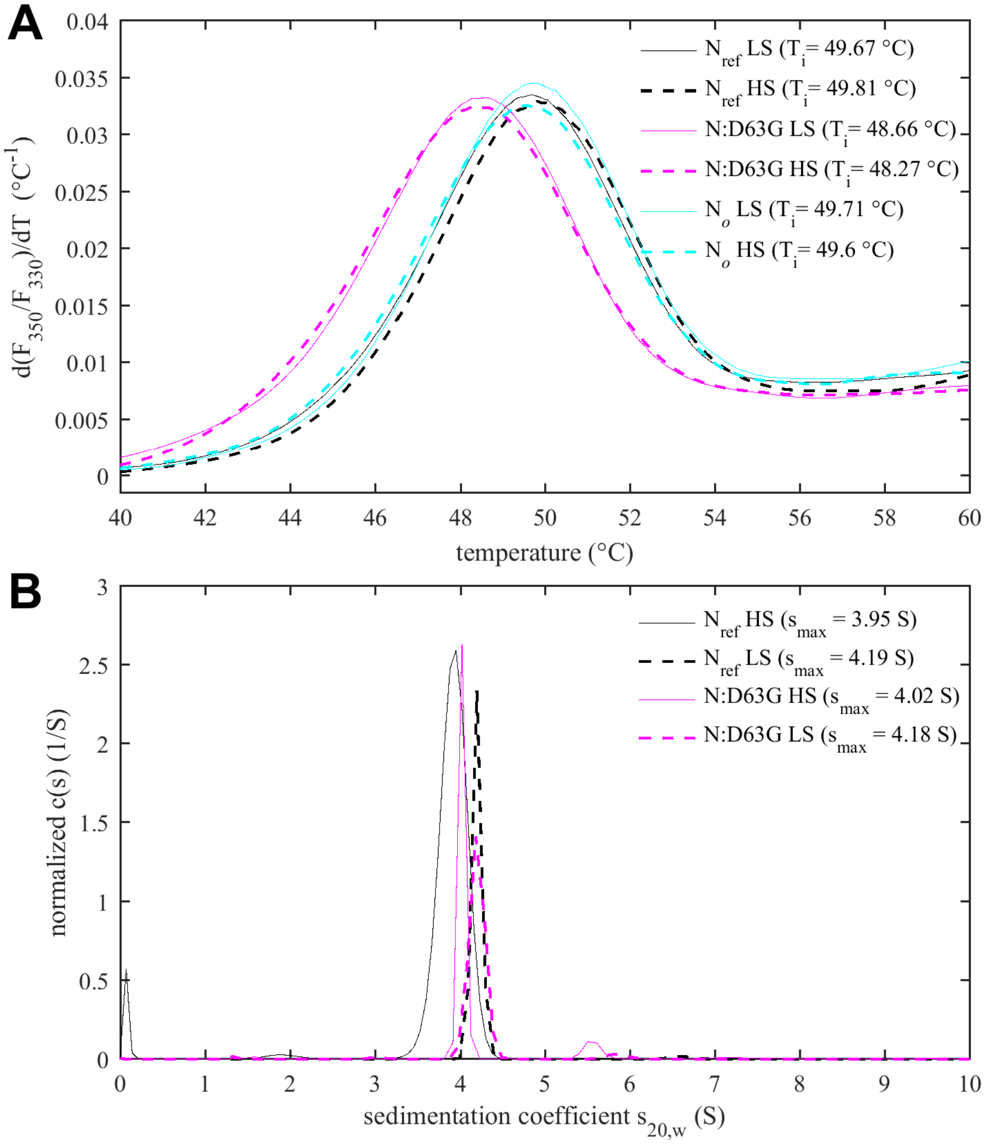


**Figure S7. Comparison of solution state of N-protein species in low-salt and high-salt buffer.** Protein preparations were dialyzed in either high-salt buffer (HS) consisting of 20mM HEPES, 150mM NaCl, pH 7.5, or low-salt buffer (LS) consisting of 10.1 mM $Na_2PO_4$, 1.8 mM $KH_2PO_4$, 2.7 mM KCl, 10 mM NaCl, pH 7.4. (**A**) DSF experiments show no significant shift in $T_i$ for the same protein species in LS or HS buffer. (**B**) SV-AUC exhibit sedimentation coefficient distributions with peak *s*-values increased by ≈5% in LS buffer relative to HS buffer. This apparent change is negligible compared to the ≈60-90% increase in sedimentation coefficients from altered oligomeric states observed for N:G215C and Nδ (**Figure 5A**).
