## Supplemental Figure S1 for "Modulation of Biophysical Properties of Nucleocapsid Protein in the Mutant Spectrum of SARS-CoV-2"

**Supplementary Figure S1:**

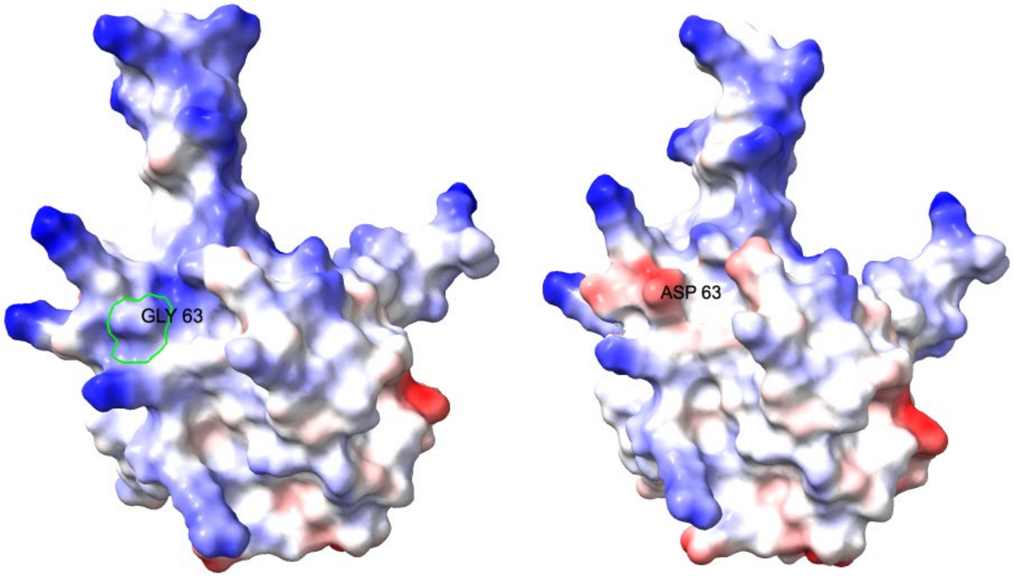


**Figure S1. Structural comparison of N:D63G mutant and ancestral N-protein.** Structures are predicted using ColabFold for N:D63G (left) and the ancestral protein (right).
