## Supplemental Figure S2 for "Modulation of Biophysical Properties of Nucleocapsid Protein in the Mutant Spectrum of SARS-CoV-2"

**Supplementary Figure S2:**

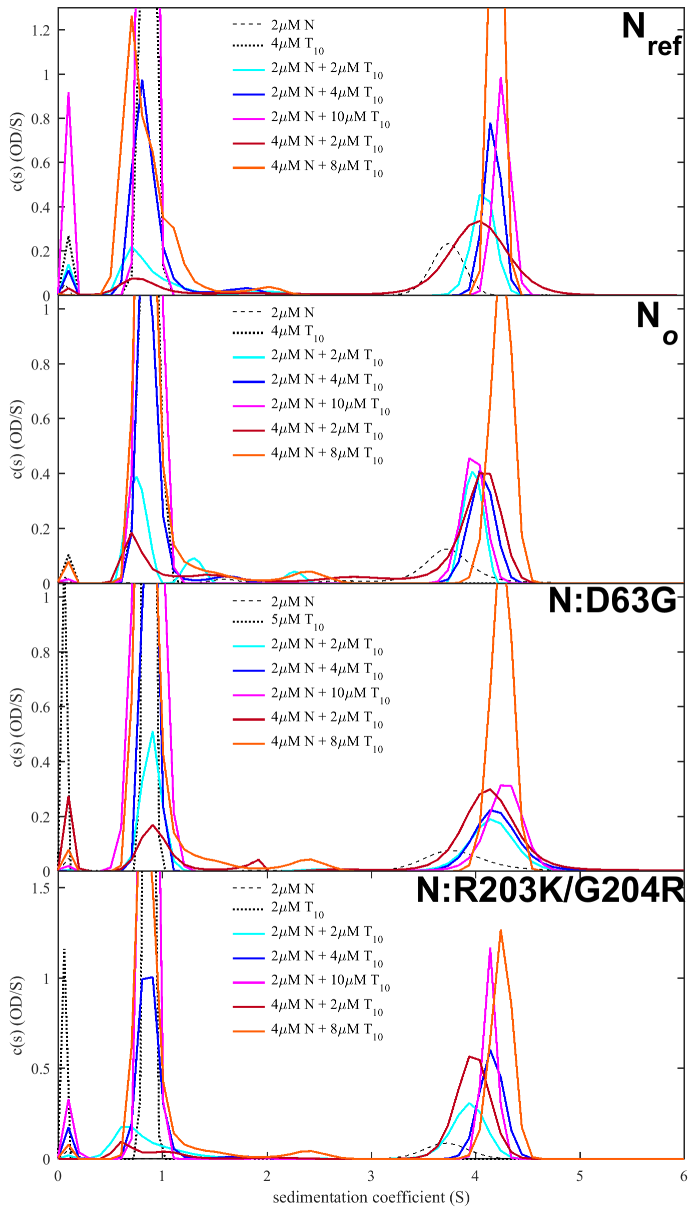


**Figure S2. N-protein affinity for binding nucleic acids probed by SV-AUC of N-protein mixtures with oligonucleotide $T_{10}$.** $T_{10}$ can occupy the NA binding grove of the NTD of N-protein, but does not permit multi-valent binding. Titration series of N-protein with $T_{10}$ allows separation of concentration-dependent populations of free and bound/co-migrating $T_{10}$ in the mixtures. This provides the basis for the determination of equilibrium binding constants through non-linear regression of the isotherm of signal-weighted average sedimentation coefficients using a two-site binding model of $T_{10}$ to N-protein dimers. Best-fit $K_D$-values and 95% confidence intervals are 1.1 [0.8 – 1.6] μM for $N_{ref}$, 2.8 [1.6 – 4.9] μM for $N_o$, 2.4 [1.1 – 5.0] μM for N:D63G, and 1.3 [0.9 – 1.9] μM for N:R203K/G204R, respectively. SV-AUC experiments were carried out in buffer HS. Similarly, no significant difference was measured in binding affinity between $N_{ref}$ and N:D63G was observed in buffer LS.
